# Engineering the smallest transcription factor: accelerated evolution of a 63-amino acid peptide dual activator-repressor

**DOI:** 10.1101/725739

**Authors:** Andreas K. Brödel, Rui Rodrigues, Alfonso Jaramillo, Mark Isalan

## Abstract

Transcription factors control gene expression in all life. This raises the question of what is the smallest protein that can support such activity. In nature, Cro from bacteriophage λ is the smallest known repressor (66 amino acids; a.a.) but activators are typically much larger (e.g. λ cI, 237 a.a.). Indeed, previous efforts to engineer a minimal activator from Cro resulted in no activity *in vivo*. In this study, we show that directed evolution results in a new Cro activator-repressor that functions as efficiently as λ cI, *in vivo*. To achieve this, we develop Phagemid-Assisted Continuous Evolution: PACEmid. We find that a peptide as small as 63-a.a. functions efficiently as an activator and/or repressor. To our knowledge, this is the smallest protein gene regulator reported to date, highlighting the capacity of transcription factors to evolve from very short peptide sequences.

---

DNA-binding proteins that regulate transcription initiation and control gene expression are called transcription factors (TFs). The question of, “*what is the smallest peptide that can function as a TF?*” is a fundamental one, with broad implications for the evolution of gene regulation (*1–3*). However, the potential for a peptide to be a minimal TF depends on its function: whether it is an activator, a repressor, or has dual activity.

In bacteria, TFs usually recruit or block RNA polymerase (RNAP), in order to activate or repress genes of interest, respectively. Generally, repression is more straightforward to achieve because it only requires a DNA-binding protein to occlude key motifs in a promoter, or to block transcription elongation by ‘roadblock’ (*4*). By contrast, activator TFs have to strike a balance between DNA binding, RNA-polymerase recruitment and RNA-polymerase release, to initiate transcription efficiently (*5*). Consequently, activators and dual TFs might need to be larger and more complex than repressors. These include well-studied examples such as λ cI (237 a.a.) from bacteriophage λ’s genetic switch (*6*) (**Fig. S1A**).

In this study, we set out to test the minimal size limits for TFs, and in particular whether activators or dual TFs could be made smaller than the smallest known repressors. Viruses have some of the smallest functional TFs and, to our knowledge, phage λ Cro protein is the smallest TF (repressor) characterized to date, containing only 66 amino acids (*7, 8*). Cro controls the viral life cycle along with λ cI and together they function naturally as a toggle switch (*6*) (**Fig. S1B**).

It has been previously shown that Cro might potentially be converted into an activator by transferring the activating surface patch of λ cI onto the surface of Cro (*9*). However, this rationally-engineered Cro variant had only a trace of detectable activity *in vitro* and none *in vivo*, in *E. coli*, because of its low affinity for λ operators (*9*). Cro is significantly smaller than common transcription factors (e.g. LacI, 360 a.a.; TetR, 207 a.a.) making it an ideal scaffold for developing a new set of small or minimal activators or dual TFs (**Fig. S1B**). In addition, a set of Cro activators would complement the λ cI synthetic biology toolbox (*10*) for gene circuit engineering in bacterial cells (*11, 12*), because cI and Cro function on related operators.

To obtain a set of small transcriptional activators derived from λ Cro, we converted our phagemid-based batch selection system (*10, 13*) for accelerated continuous evolution, in a manner similar to Phage-Assisted Continuous Evolution (PACE) (*14, 15*). One difference between our system and classic PACE is that we use phagemids rather than phage in order to facilitate screening large libraries. We also prevent bystander mutations in phage genes by placing them on helper plasmids (HP) and accessory plasmids (AP), respectively (**Fig. 1A**). These plasmids are continuously replenished within fresh uninfected cells (**Fig. 1B**). The resulting phagemid evolution system, which we call PACEmid, is based on conditional M13 bacteriophage replication. The selection process takes place inside *E. coli* cells by linking the evolving transcription factor activity to restoring essential phage Gene VI expression (deleted from the HP). In this way, a transcription factor with novel properties can be selected after several cycles of reinfection (**Fig. 1B**) and the process can be automated (**Fig. 1C,D**).

**Fig. 1.**
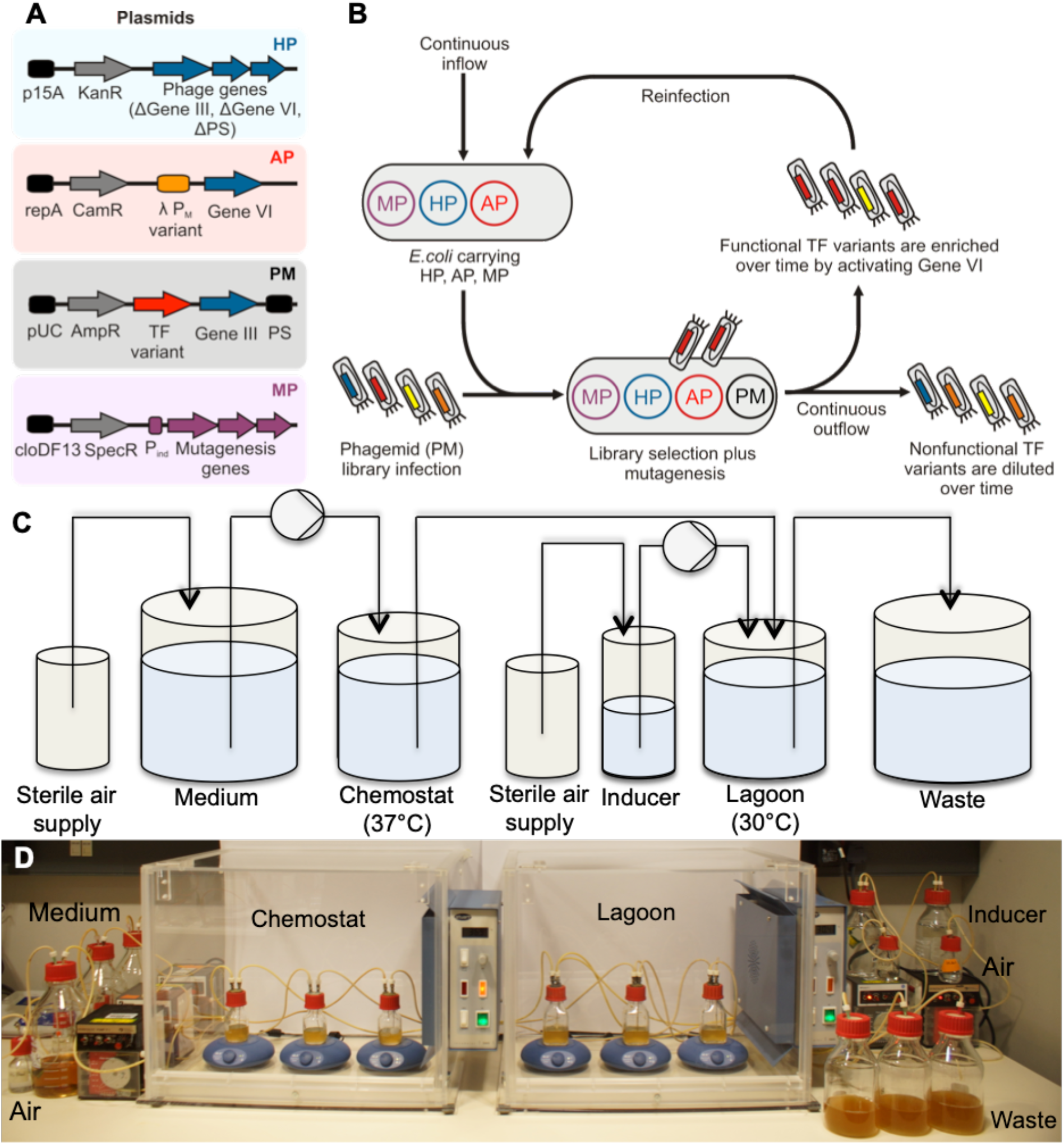
Phagemid-Assisted Continuous Evolution of transcription factors. (**A**) Plasmids: HP, Helper Phage to provide all phage genes except for gIII and gVI; AP, Accessory Plasmid to provide conditional Gene VI expression to enable selection of a successful evolving TF variant; PM, Phagemid containing an evolving TF variant and gIII; MP, chemically-inducible Mutagenesis Plasmid. (**B**) Continuous selection flow diagram: host cells (containing HP, AP, MP) get infected with M13 phage. Only an active transcription factor (TF) induces Gene VI expression to complete the phage life cycle, thus enriching this library variant; nonfunctional TF variants are diluted over time. (**C**) Flow chart of the PACEmid continuous evolution system. *E. coli* cells (containing HP, AP, MP) are cultured in the late log-phase (Chemostat, 37°C) and flow through a lagoon (30°C) containing the evolving phagemid (PM). (**D**) Photo of bioreactor setup showing three independent experiments performed in parallel.

For continuous selection, we found it essential to tune the basal Gene VI expression rate to produce sufficient amounts of phage in the absence of an active transcription factor, reducing the chances of phage loss in the lagoon. We carried out model selections with cI_opt_, (a λ cI optimized mutant with a strong activation region (*16*)) and showed that selection stringency and rate can be tuned by changing the copy number (*17*) of the accessory plasmid (AP) (**Fig. S2**). Furthermore, enrichment of cI_opt_ was more efficient in continuous mode than in batch mode, under the same selection pressure (**Fig. S3**), confirming the advantages of continuous selection (*14, 15*) for accelerated evolution.

We next implemented a mutagenesis device to expand the mutation spectrum beyond combinatorial libraries. On ColE1-derived plasmids, such as our phagemid, the leading-strand replication pol I is gradually replaced by pol III over at least 1.3kb downstream of the origin of replication (*18*). We therefore characterized the efficiency of three mutagenesis cassettes carrying error-prone pol I (EP pol I (*19*)) or error-prone pol III variants (MP4, MP6 (*20*)) under the inducible promoters P_BAD_ and P_Llac_ (*21*). Mutation rates were then analyzed by a β-lactamase stop codon reversion assay (**Fig. S4A**), a rifampicin resistance assay (**Fig. S4B,C**) and by monitoring loss of RFP function on M13 phagemids after three rounds of batch evolution (**Fig. S4D,E**). The use of the MP6 (*20*) cassette under the IPTG-inducible promoter P_Llac_ (P_Llac_-MP6-SpecR) led to the highest mutation rates and this was chosen for downstream applications (**Fig. S4F**).

To validate the use of the adapted MP6 (*20*) mutator cassette for directed evolution, we first evolved an improved orthogonal λ cI TF (cI_4A5T6T,P_; formerly the least active member of our cI toolkit (*10*)) (**Fig. S5A**). The resulting cI variant possessed a Met to Thr mutation at position 42, leading to an improved dual activation-repression of GFP and mCherry (**Fig. S5B**). The evolved cI_4A5T6T,P (T42)_ upregulated GFP expression 5.4-fold and led to a 91% repression of mCherry (**Fig. S5C,D**) (full sequences in Supplementary Materials). In comparison, expression of the parental cI_4A5T6T,P (M42)_ only displayed a 4.3-fold activation and a 58% repression. Notably, position 42 was not randomized in our previously-constructed λ cI library and thus confirmed the potential of extra mutagenesis for improved protein activities.

To see whether the optimized phagemid selection system was powerful enough to select a small Cro activator that could function in *E. coli*, we first constructed a bidirectional promoter (P_CS_/P_M,CS_), with three operator sites (O1, O2, O3) (**Fig. 2A**). O1 and O2 consisted of λ operator consensus sequence (CS), with the highest reported affinity for λ Cro (*22, 23*). As Cro binds with a very high affinity to WT O3 (*22*), a mutated O3 site was used in order to reduce potential autorepression of the P_M,CS_ promoter by λ Cro. Basal promoter activity of P_CS_/P_M,CS_, as well as the effect of Cro on expression levels, was characterized by GFP and mCherry production and showed that WT Cro repressed in both promoter directions (**Fig. 2B,C**).

**Fig. 2.**
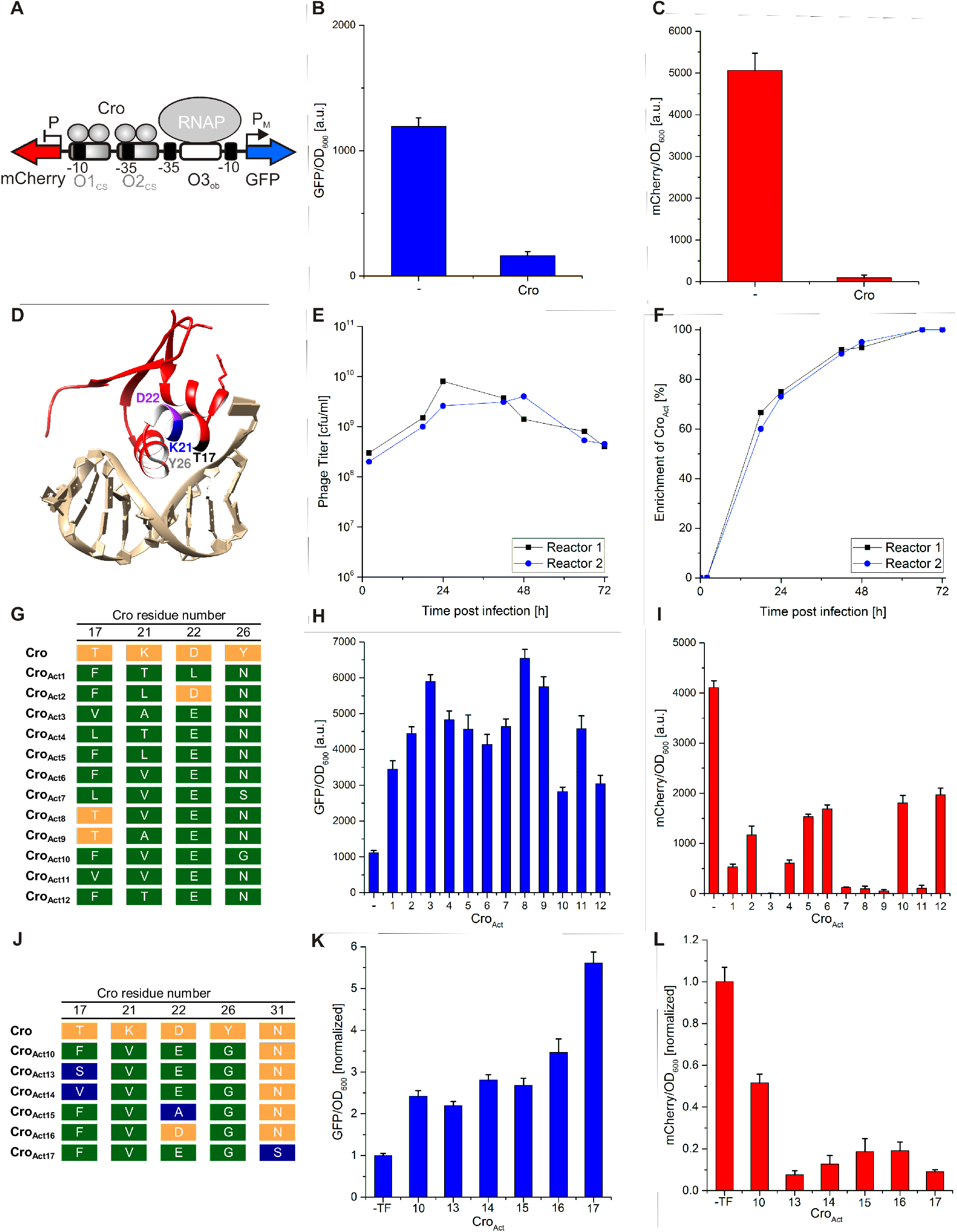
Directed evolution of Cro dual TF activator-repressors. (**A**) Bidirectional promoter designed to activate GFP and repress mCherry. Operators: O1_CS_ and O2_CS_ sites recruit Cro; O3_ob_ is designed to have weaker Cro binding. (**B**) Despite obliterating the operator O3, expression of Cro still results in partial repression of GFP under the P_M_,_CS_ promoter due to some affinity to the operator 3. (**C**) Cro binds with high affinity to the operators O1_CS_ and O2_CS_ as a dimer resulting in a strong repression of mCherry. GFP and mCherry expression was normalized to OD_600_ and data were obtained from four biological replicates. Error bars show one standard deviation. (**D**) Structural model of WT Cro binding DNA, showing key residues randomized in a combinatorial library (PDB ID: 6CRO). (**E,F**) Time-course of the phage titer and enrichment of Cro activators during selection in continuous culture, in two bioreactors. (**G**) Sequencing results of twelve selected dual Cro activators. Wild-type amino acids are highlighted in orange and non-wild-type amino acids in green. (**H,I**) Activation and repression of the bidirectional promoter P_CS_/P_M,CS_ by the selected Cro variants. GFP and mCherry expression was normalized to OD_600._ (**J**) Sequencing results from continuous directed evolution using the mutagenesis plasmid on least-active variant Cro_Act10._ (**K,L**) Activation and repression of the bidirectional promoter using the variants in (J). All data: 4 replicates; error bars are 1 standard deviation.

To search for activators, a combinatorial library of Cro variants was constructed by randomizing four amino acids in α-helix two and three (*9*) (**Fig. 2D**). Three residues (T17, K21, D22) were in the potential activation patch whereas the fourth residue (Y26) was upstream of the α-helix necessary for DNA binding. We then constructed an accessory plasmid (AP) with Gene VI under the engineered P_M,CS_ promoter, on the pSC101 vector for high selection stringency. The combinatorial Cro library was selected against this AP for three days in continuous culture leading to a reproducible enrichment of Cro activators in two separate bioreactors (**Fig. 2E,F**). The selected activators possessed at least three amino acid substitutions at the randomized positions over wild-type Cro (**Fig. 2G**). Notably, ten out of the twelve selected Cro activators contained an asparagine N at position 26. The importance of N26 for activity was confirmed with site-directed mutagenesis (**Fig. S6**). The activity of the selected Cro activators (Cro_Act_) was then analyzed with the reporter assay (**Fig. 2A)**. Cro_Act_ variants upregulated GFP expression 2.3-fold to 5.8-fold, and repressed mCherry between 52-100% (**Fig. 2H,I**). In comparison, λ cI expression resulted in a 6.7-fold activation and a 100% repression of the bidirectional P_R_/P_RM_ promoter (**Fig. S7**). The most active variants possessed the amino acids V17, A21, E22, N26 (Cro_Act3_) and T17, V21, E22, N26 (Cro_Act8_) at the randomized positions; both were strong activators and inhibitors of the dual promoter in **Fig. 2A**.

Next, we explored a wider mutation space using the mutagenesis cassette (P_Llac_-MP6-SpecR). To achieve this, we applied continuous directed evolution for four days, using an optimized pLITMUS* vector backbone (**Fig. S8**), and starting with the least-active variant Cro_Act10_ (**Fig. 2G**; **Fig. S9)**. Directed evolution was performed under a medium selection pressure (medium copy number plasmid pJPC12) compared to library selections (low copy pSC101; strong selection pressure). This resulted in higher phage production rates and thus an increased number of Cro variants. Five additional Cro activators were thus evolved with single amino acid changes in the polymerase interaction site (Cro_Act13_ to Cro_Act16_) or DNA-binding α-helix (Cro_Act17_) (**Fig. 2J-L**). Notably, a new amino acid change at position 31, N31S in the DNA-binding α-helix of Cro_Act17_, had a strong impact on the TF activity. To summarize, we obtained a set of 17 small Cro activators with a broad range of activities.

To test the potential for gene network engineering with the evolved Cro activators, we selected Cro_Act3_ (5.3-fold activation, 100% repression) for use in combination with our set of orthogonal cI variants (*10*). Successful combination would allow the construction of a wide range of gene networks that integrate multiple inputs for activation and/or repression, based on variants of commonly-used λ promoters. First, we verified the lack of cross-reactivity of the Cro_Act3,_ to any of the synthetic promoters of the orthogonal cI toolkit (**Fig. S10**). Having found no unwanted cross-reactivity, we built two different gene circuits. In the first network, gradual addition of arabinose resulted in expression of Cro_Act3_ and thus a concentration-dependent increase of GFP and a decrease of mCherry, as expected (**Fig. 3A,B**). In the second circuit, expression of the two orthogonal transcription factors Cro_Act3_ and cI_5G6G,P_, linked to the inducers arabinose and 3OC6-HSL, resulted in a concentration-dependent increase of the reporters GFP and mCherry, as designed (**Fig. 3C,D**). Therefore Cro_Act3_ can be used for gene circuit engineering, either alone or in combination with other orthogonal TFs, and can target unidirectional as well as bidirectional promoters, in a concentration-dependent manner.

**Fig. 3.**
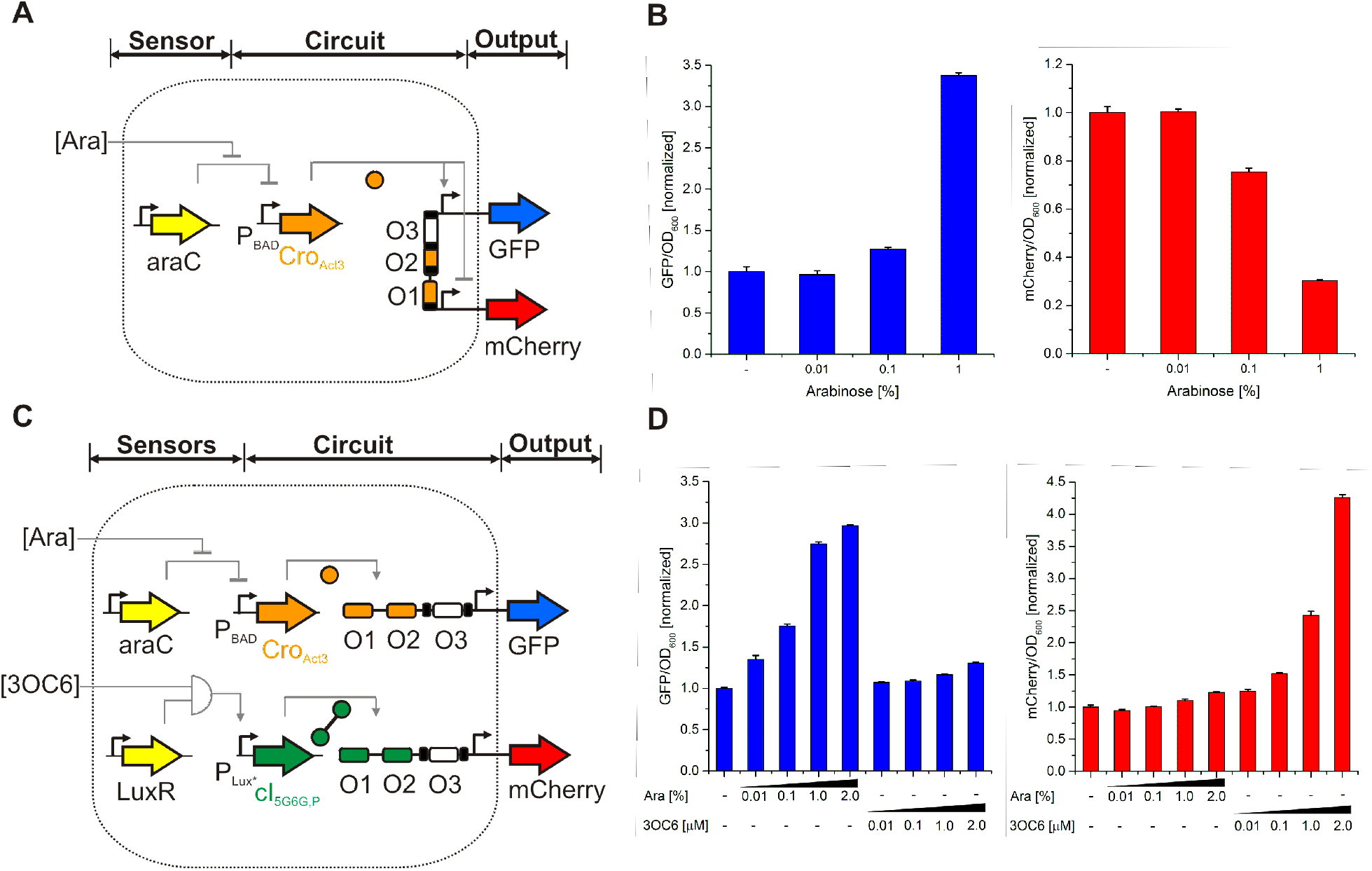
Synthetic gene circuits based on the evolved minimal activator Cro_Act3_. (**A**) Design of a 1-input gene network on a bidirectional promoter. The arabinose-inducible sensor induces Cro_Act3_ operating on a bidirectional promoter and two reporter genes. (**B**) Experimental data for the 1-input system showing the concentration-dependent response of GFP and mCherry. (**C**) Design of a 2-input gene network on two unidirectional promoters. Two sensors (P_BAD_ and P_Lux*_) act on an integrating circuit with two orthogonal TFs (Cro_Act3_ and cI_5G6G,P_) operating on two unidirectional promoters and two reporter genes. (**D**) Experimental data for the 2-input system illustrating the concentration-dependent response of GFP and mCherry to the inducers (Ara, 3OC6). All data: 3 replicates; error bars are 1 standard deviation.

At 66 a.a., Cro_Act3_ had a claim to being the smallest activator and dual TF described. However, we sought to push the boundaries of the smallest possible transcription factor. By identifying functional breakpoints in the expected structure (PDB ID: 6CRO), we made targeted deletions to the C-terminal end of the TF and analyzed the variants using our reporter assay (**Fig. 4A**). Thus, we found that a minimal 63 a.a. protein (Cro_Act3 63aa_) is still capable of transcriptional activation or repression in *E. coli* (4.4-fold activation, 88% repression) (**Fig. 4B,C**). To our knowledge, this makes it the smallest dual transcriptional factor reported to date.

**Fig. 4.**
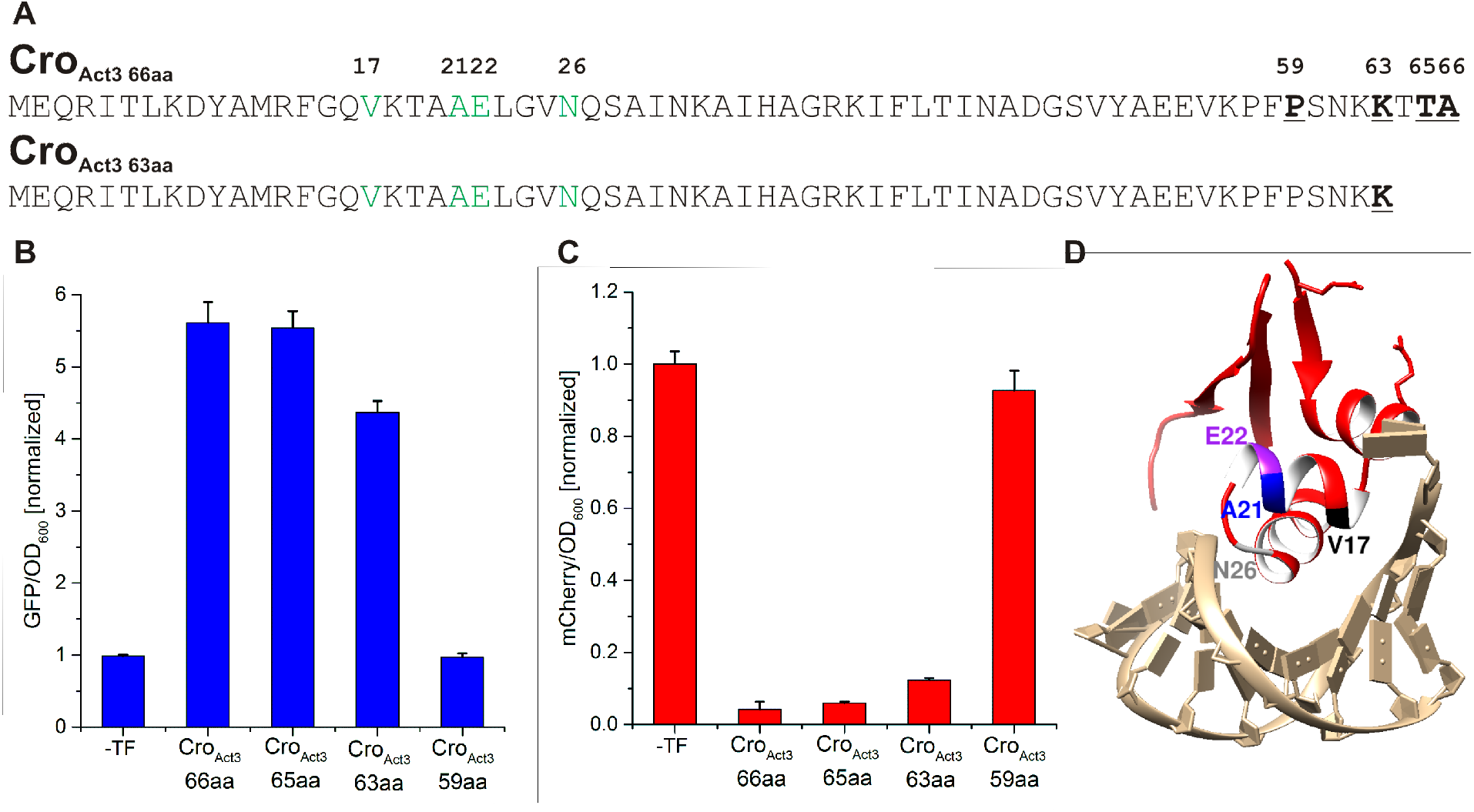
Engineering the smallest dual transcription factor. (**A**) Functional breakpoints were identified in the sequence of Cro_Act3_, including a potential structure-breaking Pro-59, a positive charge patch ending at Lys-63, and a potentially neutral C-terminal Ala-66 (bold). The corresponding truncation mutants (59, 63, 65 a.a.) were generated; Cro_Act3 63aa_ is shown as an example. Activator mutations to wild-type λ Cro repressor are highlighted in green. (**B,C**) Activation and repression of the bidirectional promoter P_CS_/P_M,CS_ by truncated Cro_Act3_ variants. GFP and mCherry expression were normalized to OD_600_; four replicates; error bars show one standard deviation. Activation and repression were normalized to the basal expression of each promoter in the absence of any TF on the phagemid. (**D**) Model indicating key features of the minimal dual TF, Cro_Act3 63aa._

We chose to evolve λ Cro because of its small size, its biological function as counterpart to λ cI, and its use in numerous synthetic biology projects (*24–27*). The evolved Cro activators described in this study can easily be constructed by site-directed mutagenesis of wild-type λ Cro, providing a straightforward approach for users to implement these Cro variants into their synthetic biology projects (components are provided in **Figs. S11-S14 and Tables S1-S7**).

One basic scientific question that we address in this study is whether a small protein repressor can be converted into a transcriptional activator *in vivo*. Overall, we conclude that not only is this possible, but that even 63 a.a. are sufficient to carry out dual intracellular transcriptional activation as well as repression. This raises interesting questions for the *de novo* evolution of gene regulation. In fully random 20 a.a. protein space, 20^63^ results in ~9 × 10^81^ combinations - an astronomically-large number for evolution to explore. However, Cro belongs to the helix-turn-helix (HTH) superfamily (*28*) and this study implies that a relatively small amount of secondary structure, including three short a-helices and three b-strands, is sufficient to make a compact scaffold that could support minimal gene regulation. As long as there are many ways to reach similar folds, the subsequent number of mutations to make a TF may be more tractable for natural selection (~11 mutations for DNA binding (*29*) and ~2-5 more for transcription activation). This implies that short peptides may have a greater capacity to evolve to regulate genes than previously thought.

## Materials and Methods

### Strains and media

Standard DNA cloning was performed with chemically competent TOP10 cells (Invitrogen) and electrocompetent TG1 cells. Combinatorial library cloning was performed with NEB 5-alpha electrocompetent cells. Phage production was carried out with TOP10, S1030 (*30*), TG1, or BL21(DE3) cells. All phage-assisted infection assays and reporter assays were performed with TG1 cells. Genotypes of all strains are listed in **Supplementary Table 1**. Cells were grown in 2×TY medium (5 g l^−1^ NaCl, 10 g l^−1^ yeast extract, 16 g l^−1^ tryptone), M9 minimal medium (6.8 g l^−1^ Na_2_HPO_4_, 3.0 g l^−1^ KH_2_PO_4_, 0.5 g l^−1^ NaCl, 1.0 g l^−1^ NH_4_Cl, 2 mM MgSO4, 100 μM CaCl_2_, 0.2% (w/v) glucose, 1 mM thiamine-HCl), or S.O.C. medium (Sigma-Aldrich). Chloramphenicol (10-25 μg ml^−1^), kanamycin (25-50 μg ml^−1^), spectinomycin (25-50 μg ml^−1^), ampicillin (50-100 μg ml^−1^), tetracycline (5-10 μg ml^−1^), and carbenicillin (10 μg ml^−1^) were added where appropriate. Isopropyl β-D-1-thiogalactopyranoside (IPTG), D-glucose or L-arabinose were added to the media to induce or repress the promoters P_Llac_ (*21*) or P_BAD_.

### Cloning and plasmid construction

Subcloning was carried out using Gibson Assembly (*31*). Green fluorescent protein (GFP, GenBank no. KM229386), red fluorescent protein (RFP), and mCherry (Uniprot no. X5DSL3) were used as reporters. The λ Cro regulatory protein (Uniprot no. P03040) was used as transcription factor scaffold and the rpoN promoter (P_rpoN_) was used to express the evolving gene on the phagemid. The stronger activator cI_opt_ (*16*) contains three amino acid changes in the λ cI gene (GenBank no., X00166) at position 35-39 (<u>S</u>VA<u>DK</u> to <u>L</u>VA<u>YE</u>). The change in copy number of accessory plasmids (pSC101, pJPC12, pJPC13) and the Cro_Act3,Y26_ variant were obtained by site-directed mutagenesis. All mutagenesis plasmids (MP4 and MP6 (*20*); EP pol I and WT pol I (*19*)) were obtained from Addgene and recloned into a vector backbone carrying a spectinomycin resistance gene (SpecR) and a cloDF13 origin of replication to make it compatible with the other plasmids of the directed evolution system. The AraC-P_BAD_ cassette on MP6-SpecR was replaced with the IPTG-inducible promoter P_Llac_ (*21*) (P_Llac_-MP6-SpecR). For the construction of gene circuits, a modified version of the P_Lux_ promoter (P_Lux*_) (*12*) was used to reduce basal expression levels. A degradation tag (AANDENYALVA) was fused to cI_5G6G,P_ at the C-terminal site to decrease basal expression in the absence of an inducer. Promoters, ribosomal binding sites and terminators were ordered as oligonucleotides (Sigma-Aldrich) or were obtained from previous studies (*32, 33*). Plasmids were purified using the QIAprep Spin Miniprep Kit or the HiSpeed Plasmid Maxi Kit (QIAGEN). Nucleotide sequences of all cloned constructs were confirmed by DNA sequencing (Eurofins Genomics). The DNA sequences of the synthetic promoters are listed in **Supplementary Fig. 11**. All plasmids and selected primer sequences are listed in **Supplementary Table 2-5**. Maps for each class of plasmid are highlighted in **Supplementary Fig. 12.**

### Construction of a combinatorial Cro library

A combinatorial Cro library was cloned based on forward and reverse primers carrying NNS codons (where S=G/C) at the positions T17, K21, D22, Y26 as described previously (*13*) (**Supplementary Table 6**). Primers were fused by PCR and fragments were cloned into the linearized pLITMUS-P_rpoN_-Cro-P_BBa_J23106_-gIII vector by Gibson Assembly. Cells were transformed and plated on 24 cm^2^ Nunc BioAssay Dishes (Thermo Scientific). Transformation efficiency was estimated by colony counting of plated serial dilutions. The next day, colonies were harvested and phagemid DNA was purified. Ten clones were sequenced to confirm diversity of the library (**Supplementary Table 7**). Molecular graphics of transcription factors were obtained with UCSF Chimera (*34*).

### Selection phage production

Selection phage production was performed in BL21(DE3) cells carrying HP-ΔPS-ΔgIII-ΔgVI and pJPC13-ΔPS-P_T7_-gVI or TOP10 cells containing HP-ΔPS-ΔgIII. Cells were made electrocompetent, phagemids transformed and cells were grown overnight at 30°C, 250 r.p.m. (Stuart Shaking Incubator SI500) in 2×TY medium supplemented with 12.5 μg ml^−1^ kanamycin, 50 μg ml ^−1^ ampicillin, and 12.5 μg ml^−1^ chloramphenicol where appropriate. For enrichment assays, plasmids carrying cI_opt_ and RFP were mixed in a ratio of 10^−6^ before transformation. 0.25 mM IPTG was added to the BL21(DE3) culture after phagemid transformation to induce Gene VI expression. Samples were centrifuged for 10 min at 8,000 g and supernatants were sterile filtered (0.22 μm pore size, Millex-GV). Phage concentration was analyzed by TG1 infection of diluted phage stocks and colony counting on ampicillin plates.

### Phagemid-assisted batch evolution

TG1 or S1030 cells carrying the helper phage HP-ΔPS-ΔgIII-ΔgVI, a Gene VI-based accessory plasmid and, optionally, a mutagenesis plasmid were grown on agar plates (M9 or LB) supplemented with appropriate antibiotics. For TG1-based evolution, starter cultures were inoculated in 2×TY with appropriate antibiotics and grown for 5-6 h at 37°C until the OD_600_ reached 0.3-0.6. For S1030-based evolution, overnight cultures were inoculated from single colonies. The next day, selection cultures were prepared with a 100-fold dilution of the overnight cultures and cells were grown for 3-4 h at 37°C until the OD_600_ reached 0.3-0.6. Cultures were infected at a desired multiplicity of infection (MOI) and a chemical inducer was added where appropriate. Cell cultures were incubated for 20h at 30°C, 250 r.p.m. (Stuart Shaking Incubator SI500). Overnight cultures were centrifuged for 10 min at 8,000 g and the phage supernatant was used to start a new round of evolution. After each round, phage supernatants were diluted, before infecting TG1 cells carrying an appropriate reporter plasmid. Infected cells were selected on ampicillin plates and single colonies were grown overnight in 2×TY medium supplemented with ampicillin. Phagemid DNA was purified using the QIAprep Spin Miniprep Kit (QIAGEN) and analyzed by sequencing (Eurofins Genomics).

### Phagemid-Assisted Continuous Evolution (PACEmid)

Glass bottles, chemostats and lagoons connected via biocompatible tubing (Cole-Parmer) were autoclaved and cooled down to room temperature. The autoclaved chemostats and lagoons were placed into two incubators (SID60, Stuart) on individual shakers (Topolino mobil, IKA). Before each experiment, the bioreactor was equilibrated by pumping 2×TY supplemented with the appropriate antibiotics through the system at 20 ml h^−1^ (Pharmacia Biotech Pump P-1). S1030 cells carrying the modified helper phage HP-ΔPS-ΔgIII-ΔgVI, an accessory plasmid and, optionally, a mutagenesis plasmid were grown on agar plates supplemented with appropriate antibiotics and 1% (w/v) D-glucose. The next day, 10 ml cultures were inoculated from single colonies, grown overnight at 37°C, 250 r.p.m. and stored at 4°C. Starter cultures were inoculated with a 100-fold dilution of the overnight culture and grown at 37°C, 250 r.p.m. until the OD_600_ reached 0.3-0.6. Chemostats were filled with 25 ml of this starter culture and cells were grown at 37°C with magnetic stir-bar agitation. The lagoon was filled with 40 ml of the starter culture and cells were infected at a MOI of four and 1mM IPTG was added where appropriate. The infected cells were grown at 30°C with magnetic stir-bar agitation. The flow-rate of 2×TY supplemented with the appropriate antibiotics was set to 20 ml h^−1^ to provide continuous supply of media. 10mM IPTG in sterile water was added to the lagoon at 2-3 ml h^−1^ to obtain a final concentration of 1mM for induced mutagenesis (Pharmacia Biotech Pump P-1). Samples were taken from the outflow of the lagoon, centrifuged and supernatants were stored at 4°C. Samples were serial diluted, before infecting TG1 cells with an appropriate reporter plasmid. Phage titers were analyzed by selection on ampicillin plates and colony counting. Single colonies were picked and grown overnight in 2×TY (100μg ml^−1^ ampicillin). Phagemid DNA was purified using the QIAprep Spin Miniprep Kit (QIAGEN) and analyzed by sequencing (Eurofins Genomics). The DNA sequences of selected TFs are listed in **Supplementary Fig. 13 and 14.** For the cI_opt_/RFP enrichment assay, TG1 cells were infected with phage dilutions and plated on ampicillin plates. The ratio of white to red colonies was analyzed by colony counting and white colonies were linked to cI_opt_ infection by colony-PCR.

### β-lactamase reversion assay

TG1 cells were transformed with the reporter plasmid pLA230 and a) cloDF-P_BAD_-polA-SpecR, b) cloDF-P_BAD_-EPpolA-SpecR, c) cloDF-P_BAD_-MP4-SpecR or d) cloDF-P_BAD_-MP6-SpecR and streaked out on LB plates supplemented with spectinomycin, kanamycin and 1% (w/v) D-glucose. The plasmid pLA230 (*19*) carries a β-lactamase gene with the ochre stop codon TAA at amino acid position 26 which is located 230 bp downstream the origin of replication. The next day, single colonies were picked and grown to the mid-log phase in the presence of 1% D-glucose. Cultures were induced with 1% (w/v) arabinose and incubated for 24h at 37°C, 250 r.p.m. (Stuart Shaking Incubator SI500). Cells were diluted and plated on LB plates in the presence or absence of ampicillin and 1% D-glucose and incubated overnight at 37°C. The ratio of the number of ampicillin resistant colonies divided by the total number of colonies on LB plates was calculated.

### Rifampicin resistance assay

TG1 cells were transformed with MP4-SpecR or MP6-SpecR and plated on LB plates with the appropriate antibiotics and 1% (w/v) D-glucose. The next day, single colonies were picked and grown to the mid-log phase in the presence of 1% D-glucose. Next, cultures were induced with 1% (w/v) arabinose and incubated for 24h at 37°C, 250 r.p.m. Cells were diluted and plated on LB plates with 1% D-glucose in the presence or absence of rifampicin and incubated in the dark for 24h at 37°C. The ratio of the number of rifampicin resistant colonies divided by the total number of colonies on LB plates was calculated.

### RFP mutation assay

S1030 cells carrying HP-ΔPS-ΔgIII-ΔgVI, pJPC12-ΔPS-P_M,CS_-RBS_BBa_B0034_-gVI and a mutagenesis plasmid (MP4-SpecR, MP6-SpecR, P_Llac_-MP6-SpecR) were grown in 2×TY medium supplemented with antibiotics and 1% D-glucose until the OD_600_ reached 0.4-0.6. Cells were infected with a RFP-carrying phagemid (pLITMUS-P_BBa_R0010_-RFP-P_BBa_J23106_-gIII) at MOI 1 and cultured in the presence or absence of an inducer (1% arabinose or 1mM IPTG) for 20h at 30°C, 250 r.p.m. Phage supernatants were harvested the following day and experiments were performed for three rounds of evolution in batch mode. TG1 cells were infected with diluted phage supernatants after each round and streaked out on LB plates supplemented with ampicillin. Plates were incubated overnight at 37°C and the following day white and red colonies were counted and the ratio calculated.

### Reporter assay

TG1 cells were transformed with a pJPC12-derived reporter plasmid and a phagemid and selected overnight on agar plates at 37°C. The next day, single colonies were picked for each biological replicate and grown 3-5 h in 1 ml 2×TY supplemented with 10 μg ml^−1^ chloramphenicol and 10 μg ml^−1^ carbenicillin at 37°C, 250 r.p.m. The cultures were diluted to OD_600_ 0.01 and 150 μl were added to each well of a 96-well plate. The absorbance at 600 nm, green fluorescence (excitation: 485 nm, emission: 520 nm) and red fluorescence (excitation: 585 nm, emission: 625 nm) were measured every 10 min in a Tecan Infinite F200 PRO plate reader (37°C, shaking between readings) until the *E. coli* cells reached stationary phase. For data analysis, fluorescence readings in the late-exponential phase (OD_600_ of 0.2 for pLITMUS, OD_600_ of 0.9 for pLITMUS*) were used. Both absorbance and fluorescence were background corrected. The fluorescence was then normalized for the number of cells by dividing by the absorbance. The average of three or four replicates and the corresponding standard deviation was calculated for each sample.

### Analysis of gene circuits

Single colonies from TG1 cells carrying a pJPC12-derived reporter plasmid and a p15A-derived TF plasmid were grown for 3-4 h in 2 ml 2×TY supplemented with 5 μg/ml chloramphenicol and 5 μg/ml carbenicillin. The cultures were diluted to OD_600_ 0.01 in a total volume of 150 μl in each well of a 96-well plate. Arabinose (0.01%, 0.1%, 1%, 2%) and N-(β-ketocaproyl)-L-homoserine lactone (3OC6-HSL; 0.01 μM, 0.1 μM, 1 μM, 2 μM) were added where appropriate. The absorbance at 600 nm, green fluorescence (excitation: 485 nm, emission: 520 nm) and red fluorescence (excitation: 585 nm, emission: 625 nm) were measured every 10 min in a Tecan Infinite F200 PRO plate reader (37°C, shaking between readings) until the *E. coli* population reached stationary phase. For data analysis, fluorescence readings in the mid to late-exponential phase were used. Both absorbance and fluorescence were background corrected. The fluorescence was then normalized for the number of cells by dividing by the absorbance. The average of three replicates and the corresponding standard deviation was calculated for each sample.

## Acknowledgments

The authors thank Daniel Blicher Holst Hansen for his support.

## Funding

This research was supported by the European Commission grant FP7-ICT-2013-10 (No. 610730, EVOPROG) and the BBSRC grant EVO-ENGINE BB/P020615/1. AJ is funded by FP7-KBBE (no 613745, PROMYS), H2020 Marie Sklodowska-Curie (no 642738, MetaRNA) and EPSRC-BBSRC (no BB/M017982/1, WISB center). MI is funded by New Investigator award no WT102944 from the Wellcome Trust U.K and by the Volkswagen Foundation.

## Author contributions

Conceived and designed the experiments: AKB, AJ, MI

Performed the experiments: AKB

Analyzed the data: AKB, MI

Contributed reagents/materials/analysis tools: RR, AJ, MI

Wrote the manuscript: AKB, MI

## Competing Interests

The authors declare no competing financial interests.

## Data and materials availability

All data is available in the manuscript or the supplementary materials.

## List of Supplementary Materials

Materials and Methods

Figures S1-S14

Tables S1-S7

References (*30–34*)

