## Supplementary Information for "Engineering the smallest transcription factor: accelerated evolution of a 63-amino acid peptide dual activator-repressor"

**This PDF file includes:**

Materials and Methods

Figs. S1 to S14

Tables S1 to S7

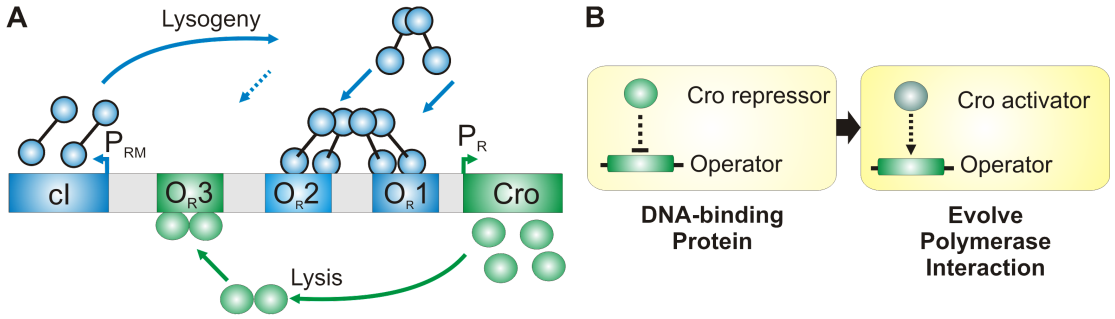


**Fig. S1. Directed evolution of small transcriptional activators based on λ Cro for biological computation in living cells.** (**A**) Schematic illustration of the phage λ switch, based on dual TF cI and repressor Cro. The $O_R$ region contains three operator sites ($O_R1$, $O_R2$, $O_R3$) and two promoters ($P_R$ and $P_{RM}$) working in opposite directions. Lambda cI has the highest affinity for $O_R1$ and $O_R2$. Binding of cI enables repression of the strong $P_R$ promoter and activation of the weak $P_{RM}$ promoter (lysogenic pathway). A very high cI concentration results in autorepression of $P_{RM}$ by binding to the $O_R3$ site (dashed line). In contrast, Cro has the highest affinity for $O_R3$ leading to repression of $P_{RM}$ (lytic pathway). To our knowledge, Cro protein is the smallest transcriptional repressor characterized to date and consists of only 66 amino acids. (**B**) Flow chart of the process to evolve and characterize small transcriptional activators based on λ Cro. There are in total six λ operators ($O_L1$, $O_L2$, $O_L3$, $O_R1$, $O_R2$, $O_R3$) from the leftward $P_L$ and the rightward $P_R$ promoters. Cro has the highest affinity to the consensus sequence (CS) of the six λ operators (*22*). The Cro protein and its consensus operator CS is used as a repressor-operator pair. Cro can potentially be turned into a transcriptional activator by evolving a polymerase interaction site into the repressor molecule. The set of small Cro activators can be used for tunable biological computation, either alone or in combination with other transcription factors.

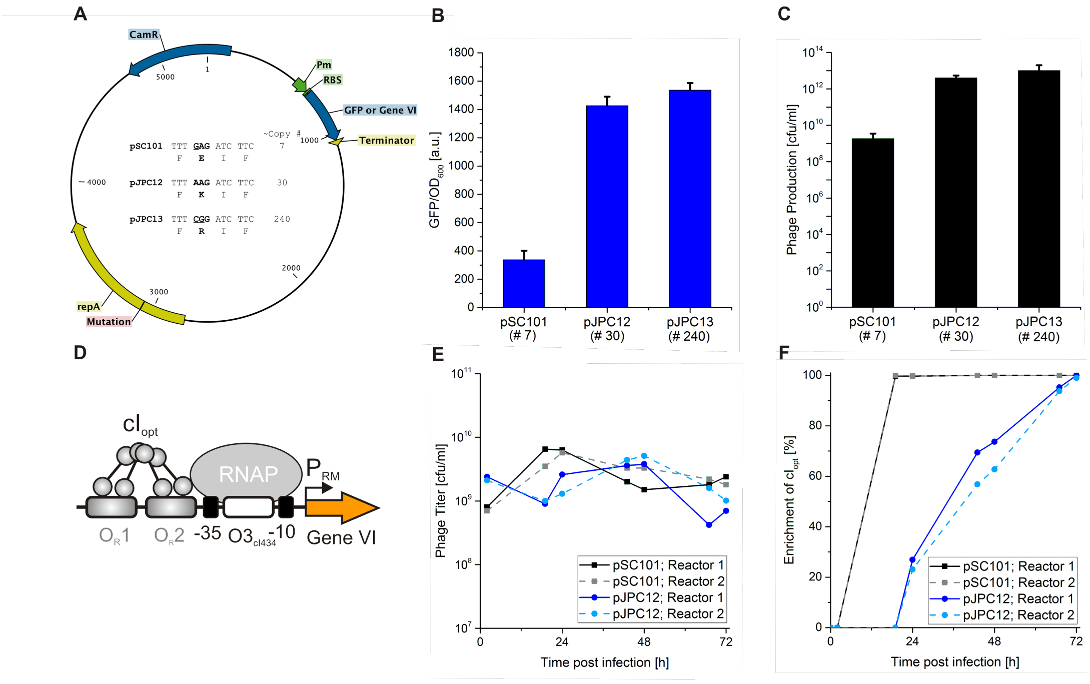


**Fig. S2. Copy number adjusts selection stringency in the Phagemid-Assisted Continuous Evolution (PACEmid) system.** (**A**) The copy number of the accessory plasmid can be modified by a single amino acid mutation in the repA origin of replication (*17*). (**B**) Effect of the copy number change on basal GFP expression under the synthetic promoter $P_{M,4A5T6T}$ (*10*). Increase of the copy number from 7 (pSC101) to 30 (pJPC12) resulted in a 4.2-fold upregulation of GFP expression in a reporter assay. An additional increase of the copy number to 240 (pJPC13) only marginally increased basal GFP expression. (**C**) Phage production can be tuned by changing the copy number of the accessory plasmid. Phage titers of TG1 cells carrying the helper phage HP-Δg3-Δg6-ΔM13 and the appropriate AP ($P_{M,4A5T6T}$-Gene VI) with three different copy numbers were analyzed. Cells were infected with RFP-expressing phagemid at a MOI of two and phage production was analyzed after 20h post-infection at 30°C in batch mode. Error bars denote the standard deviation of three biological replicates. (**D**) Scheme of the accessory plasmid with Gene VI used for selection. Lambda $cI_{opt}$ (*16*) binds to the promoter and activates Gene VI expression. (**E, F**) Enrichment of $cI_{opt}$ from a mixed phage population with $10^6$-fold excess of

RFP-expressing phagemid in continuous culture. Selections were performed under two different selection pressures by having the Gene VI circuit on a low copy (pSC101, strong pressure) or a medium copy number plasmid (pJPC12, medium pressure). For each selection stringency, two independent bioreactor experiments were performed. Samples were analyzed twice a day from the outflow of each lagoon. Enrichment of $cI_{opt}$ was analyzed by calculating the ratio of white ($cI_{opt}$) to red (RFP) colonies on agar plates.

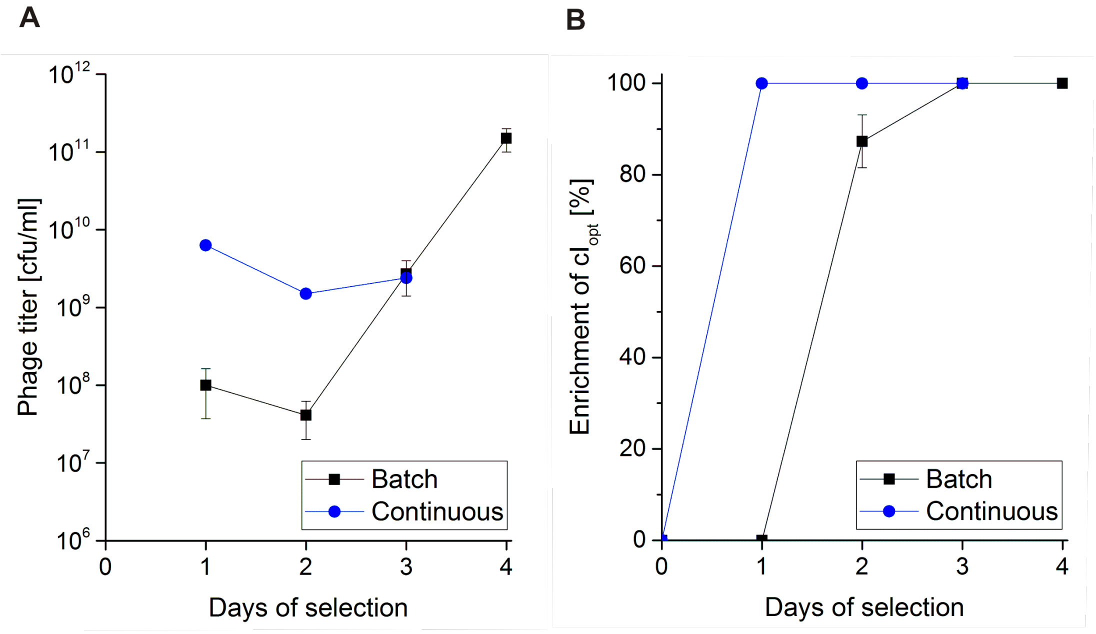


**Fig. S3. Enrichment assays of $cI_{opt}$ from mixed phage populations with $10^6$-fold excess of RFP-expressing phagemid in batch and continuous mode.** Selections were performed under the same selection pressure by having the Gene VI circuit on the low copy number plasmid pSC101 in S1030 cells. Batch cultures were infected at a multiplicity of infection of 0.1 (Round 1) and a 100-fold ratio of supernatant to cell culture was used for consecutive rounds (Round 2 to 4). (**A**) Phage titer in batch and continuous mode. For batch selections, samples were analyzed after each round. (**B**) Enrichment of $cI_{opt}$ was analyzed by calculating the ratio of white ($cI_{opt}$) to red (RFP) colonies on agar plates. In all experiments, phage encoding $cI_{opt}$ were fully enriched after the selection process. Batch data represent the average of three replicates and error bars correspond to 1 s.d. between the measurements. Continuous data show results of bioreactor 1 (see **Fig. S2 E,F**).

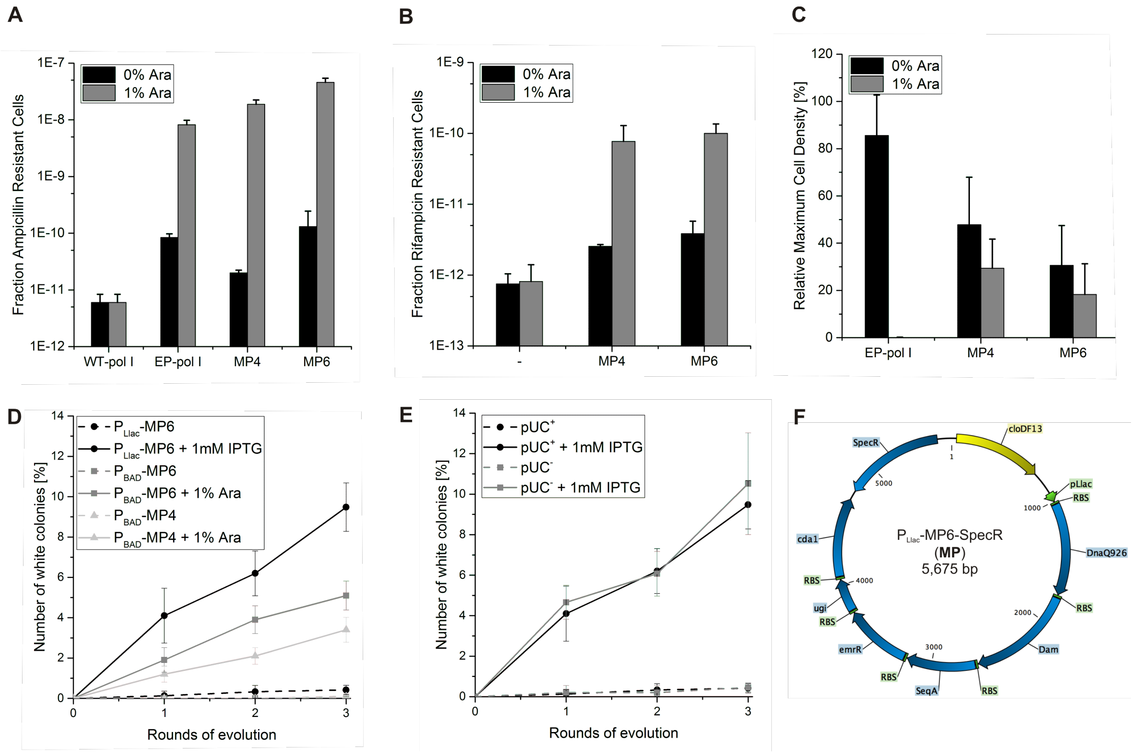


**Fig. S4. Implementation of a mutagenesis cassette into the phagemid-assisted directed evolution system**. (**A**) Ampicillin reversion assay of error-prone EP-polA, MP4, and MP6 under an arabinose-inducible promoter $P_{BAD}$ in TG1 cells. Reversion of the stop codon TAA at position 26 of the β-lactamase gene, located 230 bp downstream of the origin of replication, led to ampicillin resistance of the individual cell. WT-pol I under the $P_{BAD}$ promoter was used as a control. Mutation rates were analyzed in the presence or absence of 1% arabinose (Ara). (**B**) Rifampicin resistance assay of TG1 cells carrying MP4 or MP6 in the presence or absence of 1% arabinose. (**C**) Relative maximum cell densities in the presence or absence of 1% arabinose. Cell cultures carrying a mutagenesis plasmid were normalized to a TG1 culture. (**D**) S1030 cells carrying the HP-ΔPS-ΔgIII-ΔgVI, pJPC12-ΔPS-$P_{M,CS}$-$RBS_{BBa\_B0034}$-g6, and a mutagenesis plasmid were infected with an RFP-expressing phagemid and selected for three rounds in the presence or absence of inducer (1% arabinose or 1mM IPTG) in batch mode. Relative mutation rates were analyzed by infecting TG1 cells with diluted phage supernatants after each round of evolution prior to calculating the ratio of white (inactive RFP) to red (active RFP) colonies on ampicillin plates. (**E**) Impact of the pUC orientation on the mutation frequency. Phagemids carrying the pUC in the + or - direction were evolved for three rounds using $P_{Llac}$-MP6-SpecR and the mutation rates of expressed target protein RFP were analyzed by plate analysis. (**F**) Map of the mutagenesis plasmid with the highest mutation rate.

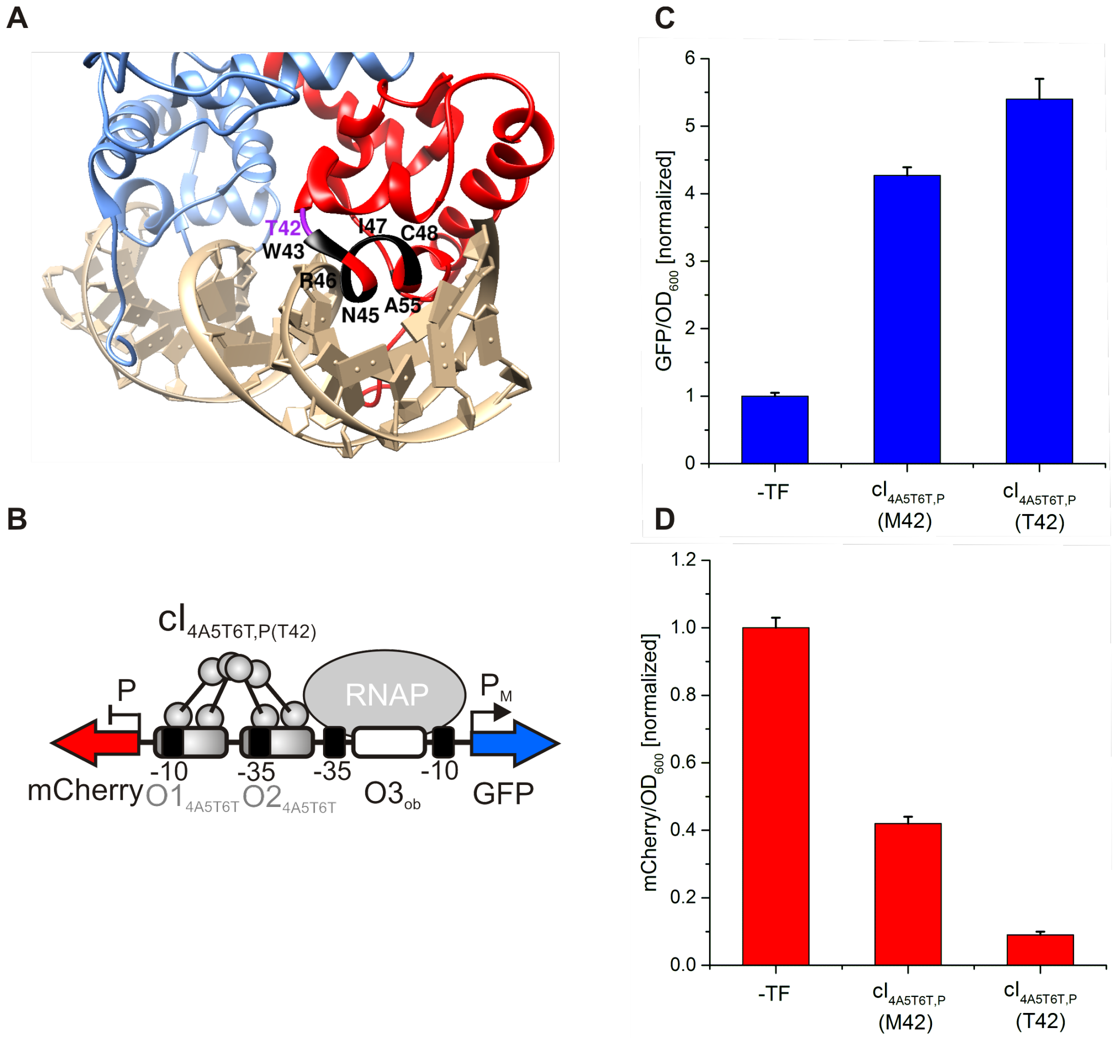


**Fig. S5. Directed evolution of an improved orthogonal cI$_{4A5T6T}$ variant using the characterized mutagenesis plasmid.** (**A**) The stronger activator of cI$_{4A5T6T,P\ (T42)}$ was evolved after five rounds of batch evolution in TG1 cells carrying the plasmids HP-ΔPS-ΔgIII-ΔgVI, pJPC12-ΔPS-P$_{M,4A5T6T}$-RBS$_{BBa_B0034}$-g6, and P$_{BAD}$-MP6-SpecR. Crystal structure of cI dimer (blue and red) binding to the operator (PDB ID: 3BDN). The amino acid changes in α-helix three of the orthogonal cI$_{4A5T6T,P}$ (*10*) variant are highlighted in black. The evolved amino acid change from M42 to T42 upstream α-helix three is highlighted in purple. (**B**) Scheme of the bidirectional promoter P/P$_{M,4A5T6T}$ used to characterize the evolved cI$_{4A5T6T,P\ (T42)}$ variant with a reporter assay. **(C, D)** The evolved cI$_{4A5T6T,P\ (T42)}$ had a Met to Thr mutation at position 42 leading to an improved DNA binding and thus an increased dual activation/ repression of GFP and mCherry compared to $cI_{4A5T6T,P\ (M42)}$. Error bars denote the standard deviation of three biological replicates. Activation and repression were normalized to the basal expression of each promoter in the absence of a transcription factor (TF) on the phagemid.

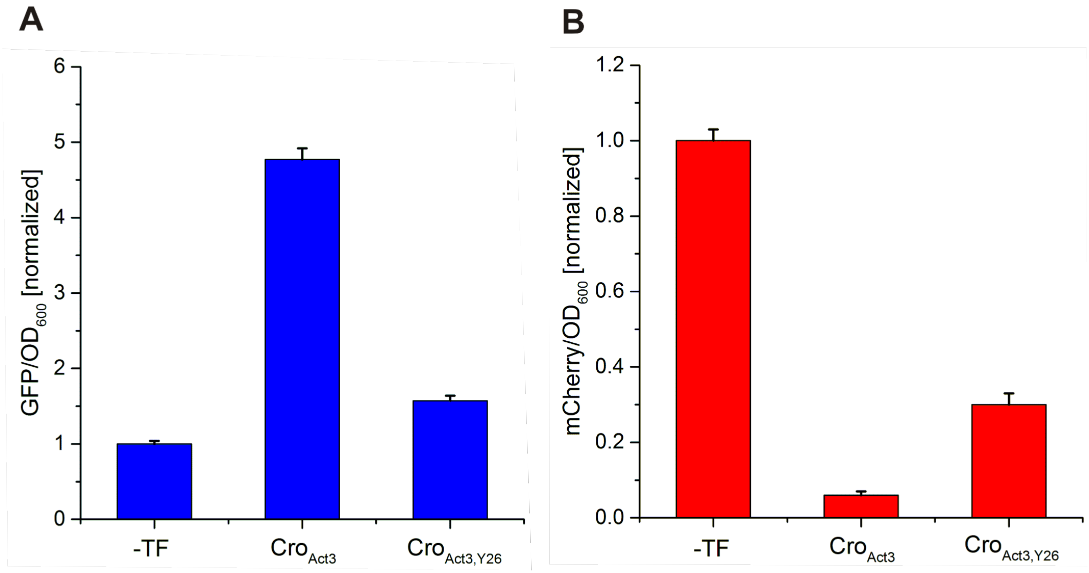


**Fig S6. Importance of asparagine at position 26 for the activity of Cro activators.** (**A**) Activation of the bidirectional promoter $P_{CS}/P_{M,CS}$ by $Cro_{Act3,Y26}$ carrying wild-type Y26 was compared to $Cro_{Act3}$ (N26) in a reporter assay. (**B**) Repression of $P_{CS}/P_{M,CS}$ by $Cro_{Act3}$ and $Cro_{Act3,Y26}$. GFP and mCherry expression was normalized to $OD_{600}$ and data were obtained from four replicates. Error bars show one standard deviation. Activation and repression were normalized to the basal expression of each promoter in the absence of a transcription factor (TF) on the phagemid.

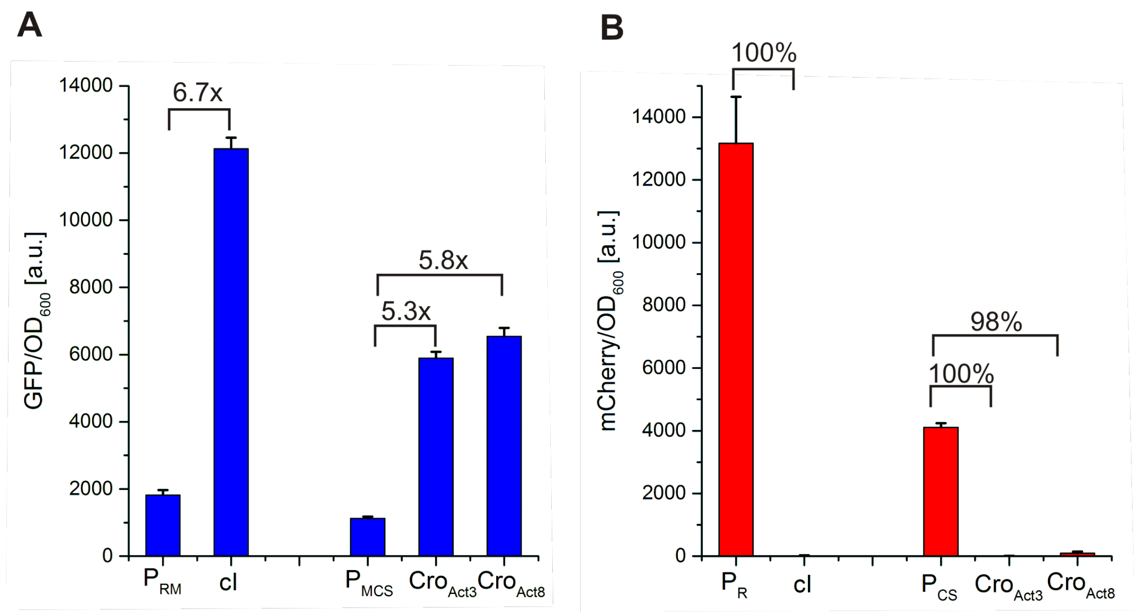


**Fig. S7. Activities of the evolved variants $Cro_{Act3}$ and $Cro_{Act8}$ compared to λ cI.** (**A, B**) The selected variants $Cro_{Act3}$ and $Cro_{Act8}$ lead to a 5.3-fold or 5.8-fold upregulation of GFP and a 100% or 98% repression of mCherry under the bidirectional promoter $P_{CS}/P_{M,CS}$. In comparison, λ cI expression resulted in a 6.7-fold activation and a full repression of mCherry under the bidirectional promoter $P_R/P_{RM}$. GFP and mCherry expression was normalized to $OD_{600}$ and data were obtained from four replicates. Error bars show one standard deviation.

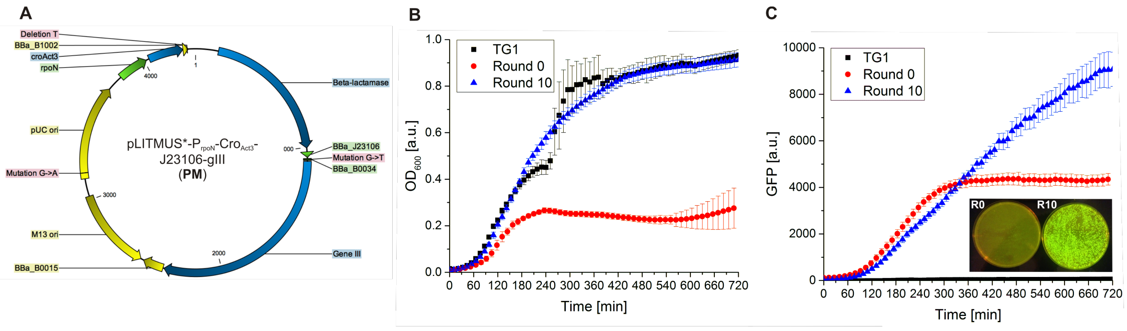


**Fig. S8. Characterization of an evolved phagemid backbone carrying $Cro_{Act3}$.** (**A**) Ten rounds of batch evolution using $P_{Llac}$-MP6-SpecR improved the phagemid backbone carrying $Cro_{Act3,}$ reducing apparent metabolic burden. The evolved phagemid contains a base pair mutation in the origin of replication, in the ribosomal binding site (RBS) upstream Gene III as well as a base pair deletion in the terminator downstream $Cro_{Act3}$. (**B**) Cell growth of TG1 cells carrying the reporter plasmid pJPC12-ΔPS-mCherry-$P_{CS}$/$P_{M,CS}$-GFP and the phagemids before or after ten rounds of batch evolution. Cells carrying the evolved phagemid (Round 10) possess an improved cell growth compared to cells with the parental phagemid (Round 0). (**C**) The improved cell growth affects the overall GFP expression of the reporter plasmid. Bacterial colonies on agar plates depict transformed *E. coli* cells under the UV light (left: Round 0; right: Round 10). Error bars denote the standard deviation of six biological replicates. TG1 cells were used a control. This improved pLITMUS* vector backbone was used in subsequent continuous evolution experiments.

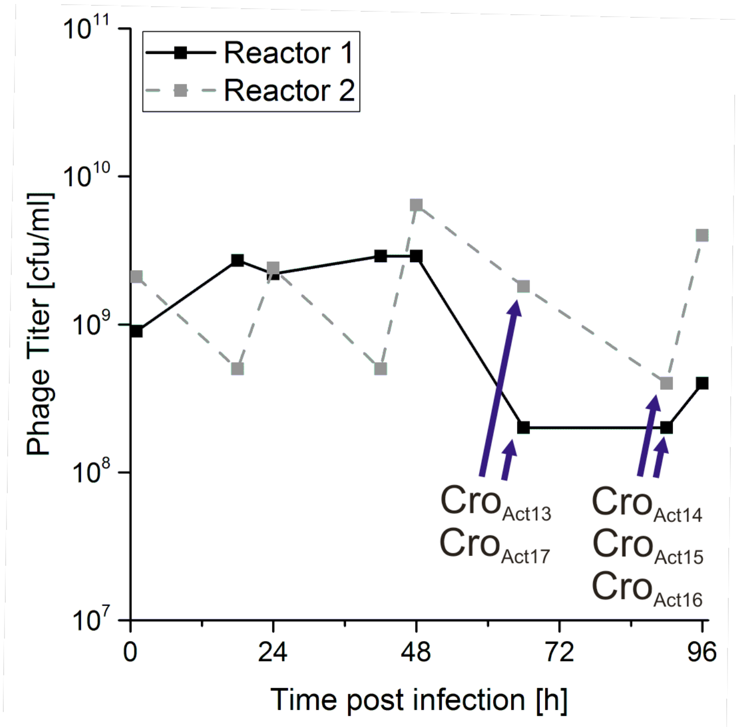


**Fig. S9. Continuous directed evolution starting from the least active variant Cro$_{Act10}$.** Phage concentrations were measured for two independent bioreactor experiments and Cro variants were obtained at the annotated time points.

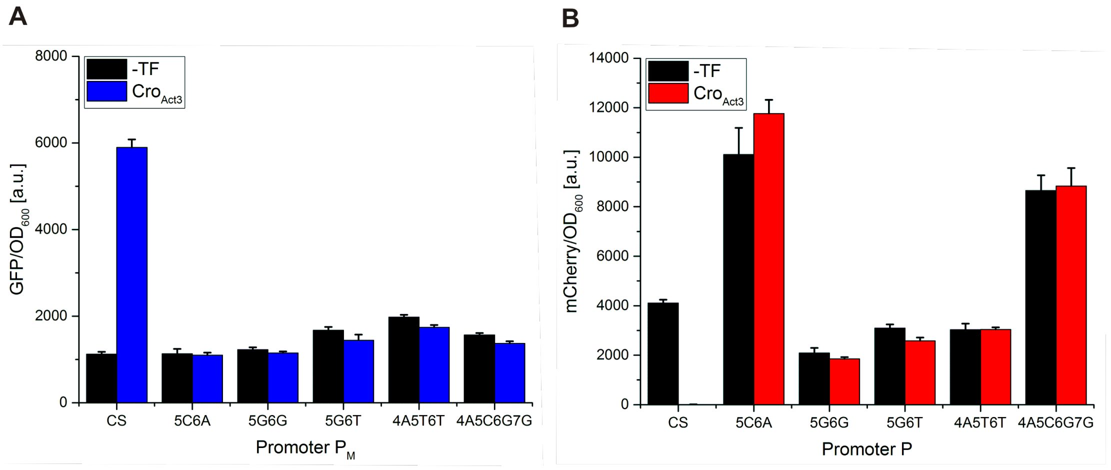


**Fig. S10. Cross-reactivity assay: the selected variant $Cro_{Act3}$ does not result in dual activation/ repression of the bidirectional promoters constructed for the orthogonal cI toolkit.** (**A**) Basal promoter strengths of the engineered promoters $P_M$ and their activation by $Cro_{Act3}$. The selected $Cro_{Act3}$ variant upregulates GFP under the consensus promoter $P_{M,CS}$ but not under any other promoter. (**B**) Basal promoter strengths of the engineered promoters P and their repression by $Cro_{Act3}$. The selected $Cro_{Act3}$ variant represses mCherry under the consensus promoter $P_{CS}$ but not under any other synthetic promoter. GFP and mCherry expression was normalized to $OD_{600}$ and data were obtained from four replicates. Error bars show one standard deviation.

O1 O2 O3

**$P_R/P_{RM}$ (wild-type)**

GCAACCATTATCACCGCCAGAGGTAAAATAGTCAACACGCACGGTGTTAGATATTTATCCCTTGCGGTGATAGATTTAACGT
CGTTGGTAATAGTGGCGGTCTCCATTTTATCAGTTGTGCGTGCCACAATCTATAAATAGGGAACGCCACTATCTAAATTGCA

**R1 R2 R3**

**$P_R/P_{RM\_Reporter}$**

GCAACCATTATCACCGCCAGAGGTAAAATAGTCAACACGCACGGTGTTAGATATTTATAAATAGTGGTGATAGATTTAACGT
CGTTGGTAATAGTGGCGGTCTCCATTTTATCAGTTGTGCGTGCCACAATCTATAAATATTTATCACCACTATCTAAATTGCA

**R1 R2 obliterated**

**$P_R/P_{RM\_Selection}$**

GCAACCATTATCACCGCCAGAGGTAAAATAGTCAACACGCACGGTGTTAGATATTACAAACTTTCTTGTATAGATTTACAAT
CGTTGGTAATAGTGGCGGTCTCCATTTTATCAGTTGTGCGTGCCACAATCTATAATGTTTGAAAGAACATATCTAAATGTTA

**R1 R2 OR1 434**

**$P_{CS}/P_{M,CS\_Reporter}$**

GCAACCATTATCACCGCCGGTGATAAAATAGTCAACACCGGCGGTGATAGATATTTATAAATAGTGGTGATAGATTTAACGT
CGTTGGTAATAGTGGCGGCCACTATTTTATCAGTTGTGGCCGCCACTATCTATAAATATTTATCACCACTATCTAAATTGCA

**CS CS obliterated**

**$P_{CS}/P_{M,CS\_Selection}$**

GCAACCATTATCACCGCCGGTGATAAAATAGTCAACACCGGCGGTGATAGATATTACAAACTTTCTTGTATAGATTTACAAT
CGTTGGTAATAGTGGCGGCCACTATTTTATCAGTTGTGGCCGCCACTATCTATAATGTTTGAAAGAACATATCTAAATGTTA

**CS CS OR1 434**

**$P/P_{M,4A5T6T\_Reporter}$**

GCAACCATTAT**ATT**CGCCG**AAT**ATAAAATAGTCAA**ATT**CGGCG**AAT**ATAGATATTTATAAATAGTGGTGATAGATTTAACGT
CGTTGGTAATA**TAA**GCGGC**TTA**TATTTTATCAGTT**TAA**GCCGC**TTA**TATCTATAAATATTTATCACCACTATCTAAATTGCA

**4A5T6T 4A5T6T obliterated**

**$P/P_{M,4A5T6T\_Selection}$**

GCAACCATTAT**ATT**CGCCG**AAT**ATAAAATAGTCAA**ATT**CGGCG**AAT**ATAGATATTTATCACCGCCGGTGATAGATTTAACGT
CGTTGGTAATA**TAA**GCGGC**TTA**TATTTTATCAGTT**TAA**GCCGC**TTA**TATCTATAAATAGTGGCGGCCACTATCTAAATTGCA

**4A5T6T 4A5T6T CS**

**Fig. S11. Sequences of synthetic promoters.** Synthetic promoters were derived from the natural bidirectional $P_R/P_{RM}$ promoter. Operators are highlighted as follows: O1 blue, O2 green, O3 grey. The natural operator O3 was modified (red) in order to bypass autorepression at high cI concentrations. For $cI_{opt}$ enrichment assays, the OR1 sequence of phage 434 was inserted at position O3. For the evolution of Cro activators, the consensus λ sequence (CS) that is based on the six natural λ operators

($O_L1$, $O_L2$, $O_L3$, $O_R1$, $O_R2$, $O_R3$) from the leftward $P_L$ and the rightward $P_R$ promoters was used at position O1 and O2 because Cro forms the most stable complex with this CS operator. WT cI binding to O3 of the orthogonal promoter $P/P_{M,4A5T6T\_Selection}$ was restored for counterselections by inserting the CS at O3.

O1 O2 O3

**P/P$_{M,5C6A}$**

```
GCAACCATTATCCACGCCGTGGATAAAATAGTCAACCACGGCGTGGATAGATATTTATAAATAGTGGTGATAGATTTAACGT
CGTTGGTAATAGGTGCGGCACCTATTTTATCAGTTGGTGCCGCACCTATCTATAAATATTTATCACCACTATCTAAATTGCA
```

**5C6A** **5C6A** **obliterated**

**P/P$_{M,5G6G}$**

```
GCAACCATTATCGGCGCCGCCGATAAAATAGTCAACGGCGGCGCCGATAGATATTTATAAATAGTGGTGATAGATTTAACGT
CGTTGGTAATAGCCGCGGCGGCTATTTTATCAGTTGCCGCCGCGGCTATCTATAAATATTTATCACCACTATCTAAATTGCA
```

**5G6G** **5G6G** **obliterated**

**P/P$_{M,5G6T}$**

```
GCAACCATTATCGTCGCCGACGATAAAATAGTCAACGTCGGCGACGATAGATATTTATAAATAGTGGTGATAGATTTAACGT
CGTTGGTAATAGCAGCGGCTGCTATTTTATCAGTTGCAGCCGCTGCTATCTATAAATATTTATCACCACTATCTAAATTGCA
```

**5G6T** **5G6T** **obliterated**

**P/P$_{M,4A5C6G7G}$**

```
GCAACCATTATACGGGCCCCGTATAAAATAGTCAAACGGGGCCCGTATAGATATTTATAAATAGTGGTGATAGATTTAACGT
CGTTGGTAATATGCCCGGGGCATATTTTATCAGTTTGCCCCGGGCATATCTATAAATATTTATCACCACTATCTAAATTGCA
```

**4A5C6G7G** **4A5C6G7G** **obliterated**

**Fig. S11 (continued). Sequences of synthetic promoters.** Synthetic promoters were derived from the natural bidirectional $P_R/P_{RM}$ promoter. Operators are highlighted as follows: O1 blue, O2 green, O3 grey. The natural operator O3 was modified (red) in order to bypass autorepression at high cI concentrations. For cI$_{opt}$ enrichment assays, the OR1 sequence of phage 434 was inserted at position O3. For the evolution of Cro activators, the consensus λ sequence (CS) that is based on the six natural λ operators ($O_L1$, $O_L2$, $O_L3$, $O_R1$, $O_R2$, $O_R3$) from the leftward $P_L$ and the rightward $P_R$ promoters was used at position O1 and O2 because Cro forms the most stable complex with this CS operator. WT cI binding to O3 of the orthogonal promoter P/P$_{M,4A5T6T_Selection}$ was restored for counterselections by inserting the CS at O3.

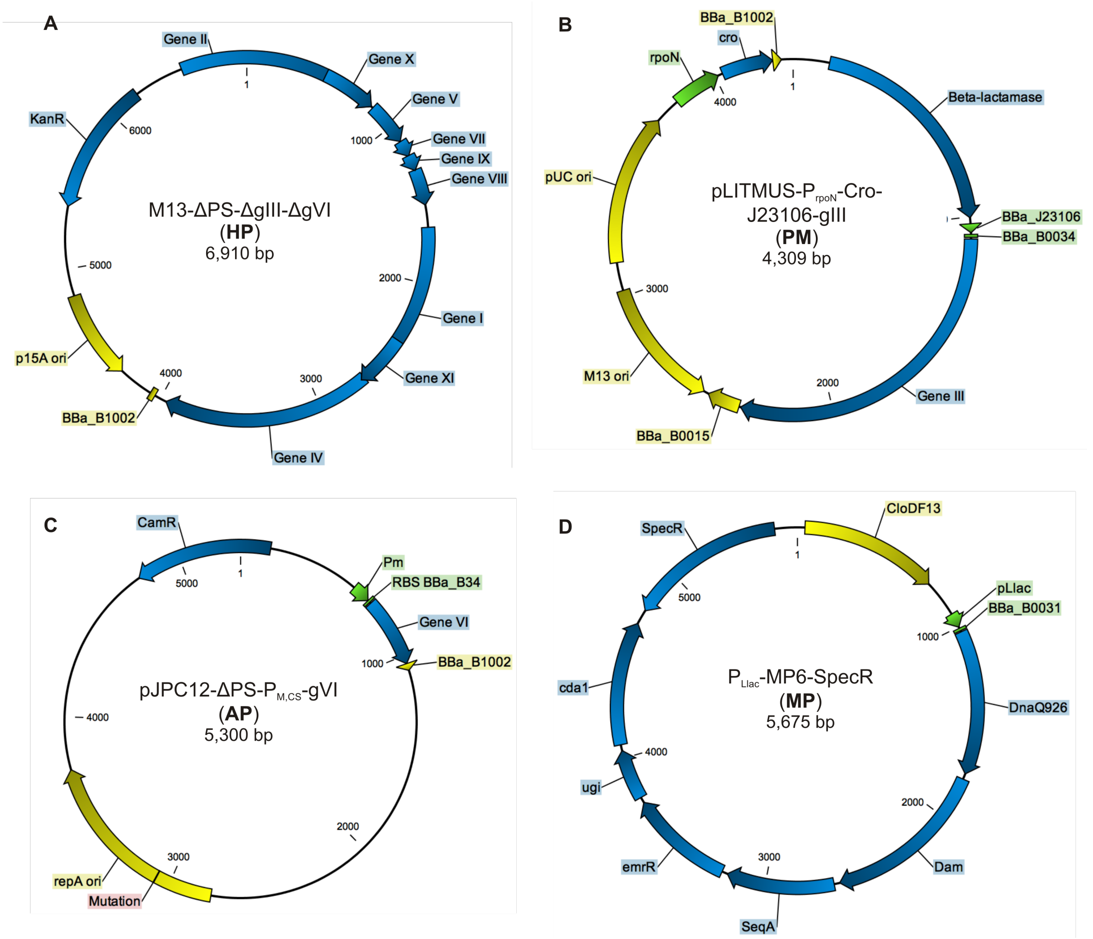


**Fig. S12. Maps of plasmids used in the Phagemid-Assisted Continuous Evolution (PACEmid) system**. (**A**) The modified helper phage M13KO7-ΔPS-ΔgeneIII-ΔgeneVI (HP) contains all phage genes required for phage replication except the genes III and VI. The weak packaging signal (PS) is removed to bypass helper phage propagation. (**B**) The phagemid pLITMUS-$P_{rpoN}$-Cro-$P_{BBa\_J23106}$-gIII (PM) provides the evolving gene of interest, the M13 packaging signal (PS) as well as constitutively expressed Gene III. (**C**) The accessory plasmid pJPC12-ΔPS-$P_{M,CS}$-$RBS_{BBa\_B0034}$-gVI (AP) contains a conditional Gene VI expression circuit, activated by an active library member on the phagemid. The copy number of the accessory plasmid can be modified by a single amino acid mutation in the repA origin of replication. (**D**) The mutagenesis plasmid (MP) carries mutator genes (dnaQ926, dam, seqA, emrR, ugi, cda1) under the IPTG-inducible promoter $P_{Llac}$ (*21*).

**Cro**

ATGGAACAACGCATAACCCTGAAAGATTATGCAATGCGCTTTGGGCAAACCAAGACA
GCTAAAGATCTCGGCGTATATCAAAGCGCGATCAACAAGGCCATTCATGCAGGCCGA
AAGATTTTTTTAACTATAAACGCTGATGGAAGCGTTTATGCGGAAGAGGTAAAGCCC
TTCCCGAGTAACAAAAAAACAACAGCATAA

MEQRITLKDYAMRFGQTKTAKDLGVYQSAINKAIHAGRKIFLTINADGSVYAEEVKP
FPSNKKTTA*

**Cro$_{Act1}$**

ATGGAACAACGCATAACCCTGAAAGATTATGCAATGCGCTTTGGGCAATTCAAGACA
GCTACGCTCCTCGGCGTAAACCAAAGCGCGATCAACAAGGCCATTCATGCAGGCCGA
AAGATTTTTTTAACTATAAACGCTGATGGAAGCGTTTATGCGGAAGAGGTAAAGCCC
TTCCCGAGTAACAAAAAAACAACAGCATAA

MEQRITLKDYAMRFGQFKTATLLGVNQSAINKAIHAGRKIFLTINADGSVYAEEVKP
FPSNKKTTA*

**Cro$_{Act2}$**

GTGGAACAACGCATAACCCTGAAAGATTATGCAATGCGCTTTGGGCAATTCAAGACA
GCTCTGGACCTCGGCGTAAACCAAAGCGCGATCAACAAGGCCATTCATGCAGGCCGA
AAGATTTTTTTAACTATAAACGCTGATGGAAGCGTTTATGCGGAAGAGGTAAAGCCC
TTCCCGAGTAACAAAAAAACAACAGCATAA

VEQRITLKDYAMRFGQFKTALDLGVNQSAINKAIHAGRKIFLTINADGSVYAEEVKP
FPSNKKTTA*

**Fig. S13. Gene sequences of WT Cro and selected Cro activator variants.** Mutations to wild-type λ Cro are highlighted in green.

**Cro$_{Act3}$**

ATGGAACAACGCATAACCCTGAAAGATTATGCAATGCGCTTTGGGCAAGTGAAGACA
GCTGCGGAGCTCGGCGTAAACCAAAGCGCGATCAACAAGGCCATTCATGCAGGCCGA
AAGATTTTTTTAACTATAAACGCTGATGGAAGCGTTTATGCGGAAGAGGTAAAGCCC
TTCCCGAGTAACAAAAAAACAACAGCATAA

MEQRITLKDYAMRFGQVKTAAELGVNQSAINKAIHAGRKIFLTINADGSVYAEEVKP
FPSNKKTTA*

**Cro$_{Act3\ 59aa}$**

ATGGAACAACGCATAACCCTGAAAGATTATGCAATGCGCTTTGGGCAAGTGAAGACA
GCTGCGGAGCTCGGCGTAAACCAAAGCGCGATCAACAAGGCCATTCATGCAGGCCGA
AAGATTTTTTTAACTATAAACGCTGATGGAAGCGTTTATGCGGAAGAGGTAAAGCCC
TTCCCGTAA

MEQRITLKDYAMRFGQVKTAAELGVNQSAINKAIHAGRKIFLTINADGSVYAEEVKP
FP*

**Cro$_{Act3\ 63aa}$**

ATGGAACAACGCATAACCCTGAAAGATTATGCAATGCGCTTTGGGCAAGTGAAGACA
GCTGCGGAGCTCGGCGTAAACCAAAGCGCGATCAACAAGGCCATTCATGCAGGCCGA
AAGATTTTTTTAACTATAAACGCTGATGGAAGCGTTTATGCGGAAGAGGTAAAGCCC
TTCCCGAGTAACAAAAAATAA

MEQRITLKDYAMRFGQVKTAAELGVNQSAINKAIHAGRKIFLTINADGSVYAEEVKP
FPSNKK*

**Fig. S13. (continued). Gene sequences of WT Cro and selected Cro activator variants.** Mutations to wild-type λ Cro are highlighted in green.

**Cro$_{Act3\ 65aa}$**

ATGGAACAACGCATAACCCTGAAAGATTATGCAATGCGCTTTGGGCAAGTGAAGACA
GCTGCGGAGCTCGGCGTAAACCAAAGCGCGATCAACAAGGCCATTCATGCAGGCCGA
AAGATTTTTTTAACTATAAACGCTGATGGAAGCGTTTATGCGGAAGAGGTAAAGCCC
TTCCCGAGTAACAAAAAAACAACATAA

MEQRITLKDYAMRFGQVKTAAELGVNQSAINKAIHAGRKIFLTINADGSVYAEEVKP
FPSNKKTT*

**Cro$_{Act3,Y26}$**

ATGGAACAACGCATAACCCTGAAAGATTATGCAATGCGCTTTGGGCAAGTGAAGACA
GCTGCGGAGCTCGGCGTATATCAAAGCGCGATCAACAAGGCCATTCATGCAGGCCGA
AAGATTTTTTTAACTATAAACGCTGATGGAAGCGTTTATGCGGAAGAGGTAAAGCCC
TTCCCGAGTAACAAAAAAACAACAGCATAA

MEQRITLKDYAMRFGQVKTAAELGVYQSAINKAIHAGRKIFLTINADGSVYAEEVKP
FPSNKKTTA*

**Cro$_{Act4}$**

ATGGAACAACGCATAACCCTGAAAGATTATGCAATGCGCTTTGGGCAATTGAAGACA
GCTACGGAGCTCGGCGTAAACCAAAGCGCGATCAACAAGGCCATTCATGCAGGCCGA
AAGATTTTTTTAACTATAAACGCTGATGGAAGCGTTTATGCGGAAGAGGTAAAGCCC
TTCCCGAGTAACAAAAAAACAACAGCATAA

MEQRITLKDYAMRFGQLKTATELGVNQSAINKAIHAGRKIFLTINADGSVYAEEVKP
FPSNKKTTA*

**Fig. S13. (continued). Gene sequences of WT Cro and selected Cro activator variants.** Mutations to wild-type λ Cro are highlighted in green.

**Cro$_{Act5}$**

ATGGAACAACGCATAACCCTGAAAGATTATGCAATGCGCTTTGGGCAATTCAAGACA
GCTTTGGAGCTCGGCGTAAACCAAAGCGCGATCAACAAGGCCATTCATGCAGGCCGA
AAGATTTTTTTAACTATAAACGCTGATGGAAGCGTTTATGCGGAAGAGGTAAAGCCC
TTCCCGAGTAACAAAAAAACAACAGCATAA

MEQRITLKDYAMRFGQFKTALELGVNQSAINKAIHAGRKIFLTINADGSVYAEEVKP
FPSNKKTTA*

**Cro$_{Act6}$**

ATGGAACAACGCATAACCCTGAAAGATTATGCAATGCGCTTTGGGCAATTCAAGACA
GCTGTGGAGCTCGGCGTAAACCAAAGCGCGATCAACAAGGCCATTCATGCAGGCCGA
AAGATTTTTTTAACTATAAACGCTGATGGAAGCGTTTATGCGGAAGAGGTAAAGCCC
TTCCCGAGTAACAAAAAAACAACAGCATAA

MEQRITLKDYAMRFGQFKTAVELGVNQSAINKAIHAGRKIFLTINADGSVYAEEVKP
FPSNKKTTA*

**Cro$_{Act7}$**

ATGGAACAACGCATAACCCTGAAAGATTATGCAATGCGCTTTGGGCAATTGAAGACA
GCTGTGGAGCTCGGCGTAAGCCAAAGCGCGATCAACAAGGCCATTCATGCAGGCCGA
AAGATTTTTTTAACTATAAACGCTGATGGAAGCGTTTATGCGGAAGAGGTAAAGCCC
TTCCCGAGTAACAAAAAAACAACAGCATAA

MEQRITLKDYAMRFGQLKTAVELGVSQSAINKAIHAGRKIFLTINADGSVYAEEVKP
FPSNKKTTA*

**Fig. S13. (continued). Gene sequences of WT Cro and selected Cro activator variants.** Mutations to wild-type λ Cro are highlighted in green.

**$Cro_{Act8}$**

ATGGAACAACGCATAACCCTGAAAGATTATGCAATGCGCTTTGGGCAAACGAAGACA
GCTGTGGAGCTCGGCGTAAACCAAAGCGCGATCAACAAGGCCATTCATGCAGGCCGA
AAGATTTTTTTAACTATAAACGCTGATGGAAGCGTTTATGCGGAAGAGGTAAAGCCC
TTCCCGAGTAACAAAAAAACAACAGCATAA

MEQRITLKDYAMRFGQTKTAVELGVNQSAINKAIHAGRKIFLTINADGSVYAEEVKP
FPSNKKTTA*

**$Cro_{Act9}$**

ATGGAACAACGCATAACCCTGAAAGATTATGCAATGCGCTTTGGGCAAACGAAGACA
GCTGCGGAGCTCGGCGTAAACCAAAGCGCGATCAACAAGGCCATTCATGCAGGCCGA
AAGATTTTTTTAACTATAAACGCTGATGGAAGCGTTTATGCGGAAGAGGTAAAGCCC
TTCCCGAGTAACAAAAAAACAACAGCATAA

MEQRITLKDYAMRFGQTKTAAELGVNQSAINKAIHAGRKIFLTINADGSVYAEEVKP
FPSNKKTTA*

**$Cro_{Act10}$**

ATGGAACAACGCATAACCCTGAAAGATTATGCAATGCGCTTTGGGCAATTCAAGACA
GCTGTGGAGCTCGGCGTAGGGCAAAGCGCGATCAACAAGGCCATTCATGCAGGCCGA
AAGATTTTTTTAACTATAAACGCTGATGGAAGCGTTTATGCGGAAGAGGTAAAGCCC
TTCCCGAGTAACAAAAAAACAACAGCATAA

MEQRITLKDYAMRFGQFKTAVELGVGQSAINKAIHAGRKIFLTINADGSVYAEEVKP
FPSNKKTTA*

**Fig. S13. (continued). Gene sequences of WT Cro and selected Cro activator variants.** Mutations to wild-type λ Cro are highlighted in green.

**Cro$_{Act11}$**

ATGGAACAACGCATAACCCTGAAAGATTATGCAATGCGCTTTGGGCAAGTGAAGACA
GCTGTGGAGCTCGGCGTAAACCAAAGCGCGATCAACAAGGCCATTCATGCAGGCCGA
AAGATTTTTTTAACTATAAACGCTGATGGAAGCGTTTATGCGGAAGAGGTAAAGCCC
TTCCCGAGTAACAAAAAAACAACAGCATAA

MEQRITLKDYAMRFGQVKTAVELGVNQSAINKAIHAGRKIFLTINADGSVYAEEVKP
FPSNKKTTA*

**Cro$_{Act12}$**

ATGGAACAACGCATAACCCTGAAAGATTATGCAATGCGCTTTGGGCAATTCAAGACA
GCTACCGAGCTCGGCGTAAACCAAAGCGCGATCAACAAGGCCATTCATGCAGGCCGA
AAGATTTTTTTAACTATAAACGCTGATGGAAGCGTTTATGCGGAAGAGGTAAAGCCC
TTCCCGAGTAACAAAAAAACAACAGCATAA

MEQRITLKDYAMRFGQFKTATELGVNQSAINKAIHAGRKIFLTINADGSVYAEEVKP
FPSNKKTTA*

**Cro$_{Act13}$**

ATGGAACAACGCATAACCCTGAAAGATTATGCAATGCGCTTTGGGCAATCCAAGACA
GCTGTGGAGCTCGGCGTAGGGCAAAGCGCGATCAACAAGGCCATTCATGCAGGCCGA
AAGATTTTTTTAACTATAAACGCTGATGGAAGCGTTTATGCGGAAGAGGTAAAGCCC
TTCCCGAGTAACAAAAAAACAACAGCATAA

MEQRITLKDYAMRFGQSKTAVELGVGQSAINKAIHAGRKIFLTINADGSVYAEEVKP
FPSNKKTTA*

**Fig. S13. (continued). Gene sequences of WT Cro and selected Cro activator variants.** Mutations to wild-type λ Cro are highlighted in green.

**Cro$_{Act14}$**

ATGGAACAACGCATAACCCTGAAAGATTATGCAATGCGCTTTGGGCAAGTCAAGACA
GCTGTGGAGCTCGGCGTAGGGCAAAGCGCGATCAACAAGGCCATTCATGCAGGCCGA
AAGATTTTTTTAACTATAAACGCTGATGGAAGCGTTTATGCGGAAGAGGTAAAGCCC
TTCCCGAGTAACAAAAAAACAACAGCATAA

MEQRITLKDYAMRFGQVKTAVELGVGQSAINKAIHAGRKIFLTINADGSVYAEEVKP
FPSNKKTTA*

**Cro$_{Act15}$**

ATGGAACAACGCATAACCCTGAAAGATTATGCAATGCGCTTTGGGCAATTCAAGACA
GCTGTGGCGCTCGGCGTAGGGCAAAGCGCGATCAACAAGGCCATTCATGCAGGCCGA
AAGATTTTTTTAACTATAAACGCTGATGGAAGCGTTTATGCGGAAGAGGTAAAGCCC
TTCCCGAGTAACAAAAAAACAACAGCATAA

MEQRITLKDYAMRFGQFKTAVALGVGQSAINKAIHAGRKIFLTINADGSVYAEEVKP
FPSNKKTTA*

**Cro$_{Act16}$**

ATGGAACAACGCATAACCCTGAAAGATTATGCAATGCGCTTTGGGCAATTCAAGACA
GCTGTGGATCTCGGCGTAGGGCAAAGCGCGATCAACAAGGCCATTCATGCAGGCCGA
AAGATTTTTTTAACTATAAACGCTGATGGAAGCGTTTATGCGGAAGAGGTAAAGCCC
TTCCCGAGTAACAAAAAAACAACAGCATAA

MEQRITLKDYAMRFGQFKTAVDLGVGQSAINKAIHAGRKIFLTINADGSVYAEEVKP
FPSNKKTTA*

**Fig. S13. (continued). Gene sequences of WT Cro and selected Cro activator variants.** Mutations to wild-type λ Cro are highlighted in green.

**Cro$_{Act17}$**

ATGGAACAACGCATAACCCTGAAAGATTATGCAATGCGCTTTGGGCAATTCAAGACA

GCTGTGGAGCTCGGCGTAGGGCAAAGCGCGATCAGCAAGGCCATTCATGCAGGCCGA

AAGATTTTTTTAACTATAAACGCTGATGGAAGCGTTTATGCGGAAGAGGTAAAGCCC

TTCCCGAGTAACAAAAAAACAACAGCATAA

MEQRITLKDYAMRFGQFKTAVELGVGQSAISKAIHAGRKIFLTINADGSVYAEEVKP

FPSNKKTTA*

**Fig. S13. (continued). Gene sequences of WT Cro and selected Cro activator variants.** Mutations to wild-type λ Cro are highlighted in green.

**cI**

ATGAGCACAAAAAAGAAACCATTAACACAAGAGCAGCTTGAGGACGCACGTCGCCTT
AAAGCAATTTATGAAAAAAAGAAAAATGAACTTGGCTTATCCCAGGAATCTGTCGCA
GACAAGATGGGGATGGGGCAGTCAGGCGTTGGTGCTTTATTTAATGGCATCAATGCA
TTAAATGCTTATAACGCCGCATTGCTTGCAAAAATTCTCAAAGTTAGCGTTGAAGAA
TTTAGCCCTTCAATCGCCAGAGAAATCTACGAGATGTATGAAGCGGTTAGTATGCAG
CCGTCACTTAGAAGTGAGTATGAGTACCCTGTTTTTTCTCATGTTCAGGCAGGGATG
TTCTCACCTGAGCTTAGAACCTTTACCAAAGGTGATGCGGAGAGATGGGTAAGCACA
ACCAAAAAAGCCAGTGATTCTGCATTCTGGCTTGAGGTTGAAGGTAATTCCATGACC
GCACCAACAGGCTCCAAGCCAAGCTTTCCTGACGGAATGTTAATTCTCGTTGACCCT
GAGCAGGCTGTTGAGCCAGGTGATTTCTGCATAGCCAGACTTGGGGGTGATGAGTTT
ACCTTCAAGAAACTGATCAGGGATAGCGGTCAGGTGTTTTTACAACCACTAAACCCA
CAGTACCCAATGATCCCATGCAATGAGAGTTGTTCCGTTGTGGGGAAAGTTATCGCT
AGTCAGTGGCCTGAAGAGACGTTTGGCTGA

MSTKKKPLTQEQLEDARRLKAIYEKKKNELGLSQESVADKMGMGQSGVGALFNGINA
LNAYNAALLAKILKVSVEEFSPSIAREIYEMYEAVSMQPSLRSEYEYPVFSHVQAGM
FSPELRTFTKGDAERWVSTTKKASDSAFWLEVEGNSMTAPTGSKPSFPDGMLILVDP
EQAVEPGDFCIARLGGDEFTFKKLIRDSGQVFLQPLNPQYPMIPCNESCSVVGKVIA
SQWPEETFG*

**cI$_{opt}$**

ATGAGCACAAAAAAGAAACCATTAACACAAGAGCAGCTTGAGGACGCACGTCGCCTT
AAAGCAATTTATGAAAAAAAGAAAAATGAACTTGGCTTATCCCAGGAATTGGTCGCA
TACGAGATGGGGATGGGGCAGTCAGGCGTTGGTGCTTTATTTAATGGCATCAATGCA
TTAAATGCTTATAACGCCGCATTGCTTGCAAAAATTCTCAAAGTTAGCGTTGAAGAA
TTTAGCCCTTCAATCGCCAGAGAAATCTACGAGATGTATGAAGCGGTTAGTATGCAG
CCGTCACTTAGAAGTGAGTATGAGTACCCTGTTTTTTCTCATGTTCAGGCAGGGATG
TTCTCACCTGAGCTTAGAACCTTTACCAAAGGTGATGCGGAGAGATGGGTAAGCACA
ACCAAAAAAGCCAGTGATTCTGCATTCTGGCTTGAGGTTGAAGGTAATTCCATGACC
GCACCAACAGGCTCCAAGCCAAGCTTTCCTGACGGAATGTTAATTCTCGTTGACCCT
GAGCAGGCTGTTGAGCCAGGTGATTTCTGCATAGCCAGACTTGGGGGTGATGAGTTT
ACCTTCAAGAAACTGATCAGGGATAGCGGTCAGGTGTTTTTACAACCACTAAACCCA
CAGTACCCAATGATCCCATGCAATGAGAGTTGTTCCGTTGTGGGGAAAGTTATCGCT
AGTCAGTGGCCTGAAGAGACGTTTGGCTGA

MSTKKKPLTQEQLEDARRLKAIYEKKKNELGLSQELVAYEMGMGQSGVGALFNGINA
LNAYNAALLAKILKVSVEEFSPSIAREIYEMYEAVSMQPSLRSEYEYPVFSHVQAGM
FSPELRTFTKGDAERWVSTTKKASDSAFWLEVEGNSMTAPTGSKPSFPDGMLILVDP
EQAVEPGDFCIARLGGDEFTFKKLIRDSGQVFLQPLNPQYPMIPCNESCSVVGKVIA
SQWPEETFG*

**Fig. S14. Gene sequences of cI variants.** Mutations in the DNA-binding site of λ cI to obtain new binding affinities are highlighted in green whereas base pair substitutions to obtain stronger transcriptional activators are highlighted in blue.

**cI$_{4A5T6T,P}$ (M42)**

ATGAGCACAAAAAAGAAACCATTAACACAAGAGCAGCTTGAGGACGCACGTCGCCTT
AAAGCAATTTATGAAAAAAAGAAAAATGAACTTGGCTTATCCCAGGAATTGGTCGCA
TACGAGATGGGGATGTGGCAGAACCGCATCTGCGCTTTATTTAATGGCATCGCGGCA
TTAAATGCTTATAACGCCGCATTGCTTGCAAAAATTCTCAAAGTTAGCGTTGAAGAA
TTTAGCCCTTCAATCGCCAGAGAAATCTACGAGATGTATGAAGCGGTTAGTATGCAG
CCGTCACTTAGAAGTGAGTATGAGTACCCTGTTTTTTCTCATGTTCAGGCAGGGATG
TTCTCACCTGAGCTTAGAACCTTTACCAAAGGTGATGCGGAGAGATGGGTAAGCACA
ACCAAAAAAGCCAGTGATTCTGCATTCTGGCTTGAGGTTGAAGGTAATTCCATGACC
GCACCAACAGGCTCCAAGCCAAGCTTTCCTGACGGAATGTTAATTCTCGTTGACCCT
GAGCAGGCTGTTGAGCCAGGTGATTTCTGCATAGCCAGACTTGGGGGTGATGAGTTT
ACCTTCAAGAAACTGATCAGGGATAGCGGTCAGGTGTTTTTACAACCACTAAACCCA
CAGTACCCAATGATCCCATGCAATGAGAGTTGTTCCGTTGTGGGGAAAGTTATCGCT
AGTCAGTGGCCTGAAGAAACGTTTGGCTGA

MSTKKKPLTQEQLEDARRLKAIYEKKKNELGLSQELVAYEMGMWQNRICALFNGIAA
LNAYNAALLAKILKVSVEEFSPSIAREIYEMYEAVSMQPSLRSEYEYPVFSHVQAGM
FSPELRTFTKGDAERWVSTTKKASDSAFWLEVEGNSMTAPTGSKPSFPDGMLILVDP
EQAVEPGDFCIARLGGDEFTFKKLIRDSGQVFLQPLNPQYPMIPCNESCSVVGKVIA
SQWPEETFG*

**cI$_{4A5T6T,P}$ (T42)**

ATGAGCACAAAAAAGAAACCATTAACACAAGAGCAGCTTGAGGACGCACGTCGCCTT
AAAGCAATTTATGAAAAAAAGAAAAATGAACTTGGCTTATCCCAGGAATTGGTCGCA
TACGAGATGGGGACGTGGCAGAACCGCATCTGCGCTTTATTTAATGGCATCGCGGCA
TTAAATGCTTATAACGCCGCATTGCTTGCAAAAATTCTCAAAGTTAGCGTTGAAGAA
TTTAGCCCTTCAATCGCCAGAGAAATCTACGAGATGTATGAAGCGGTTAGTATGCAG
CCGTCACTTAGAAGTGAGTATGAGTACCCTGTTTTTTCTCATGTTCAGGCAGGGATG
TTCTCACCTGAGCTTAGAACCTTTACCAAAGGTGATGCGGAGAGATGGGTAAGCACA
ACCAAAAAAGCCAGTGATTCTGCATTCTGGCTTGAGGTTGAAGGTAATTCCATGACC
GCACCAACAGGCTCCAAGCCAAGCTTTCCTGACGGAATGTTAATTCTCGTTGACCCT
GAGCAGGCTGTTGAGCCAGGTGATTTCTGCATAGCCAGACTTGGGGGTGATGAGTTT
ACCTTCAAGAAACTGATCAGGGATAGCGGTCAGGTGTTTTTACAACCACTAAACCCA
CAGTACCCAATGATCCCATGCAATGAGAGTTGTTCCGTTGTGGGGAAAGTTATCGCT
AGTCAGTGGCCTGAAGAAACGTTTGGCTGA

MSTKKKPLTQEQLEDARRLKAIYEKKKNELGLSQELVAYEMGTWQNRICALFNGIAA
LNAYNAALLAKILKVSVEEFSPSIAREIYEMYEAVSMQPSLRSEYEYPVFSHVQAGM
FSPELRTFTKGDAERWVSTTKKASDSAFWLEVEGNSMTAPTGSKPSFPDGMLILVDP
EQAVEPGDFCIARLGGDEFTFKKLIRDSGQVFLQPLNPQYPMIPCNESCSVVGKVIA
SQWPEETFG*

**Fig. S14. (continued). Gene sequences of cI variants.** Mutations in the DNA-binding site of λ cI to obtain new binding affinities are highlighted in green whereas base pair substitutions to obtain stronger transcriptional activators are highlighted in blue.
